# Structure-function analysis of ZAR1 immune receptor reveals key molecular interactions for activity

**DOI:** 10.1101/592824

**Authors:** Maël Baudin, Karl J. Schreiber, Eliza C. Martin, Andrei J. Petrescu, Jennifer D. Lewis

**Author notes:** These authors contributed equally (Karl J. Schreiber, Eliza C. Martin).

## Abstract

NLR (Nucleotide-binding [NB] Leucine-rich repeat [LRR] Receptor) proteins are critical for inducing immune responses in response to pathogen proteins, and must be tightly regulated to prevent spurious activation in the absence of a pathogen. The ZAR1 NLR recognizes diverse effector proteins from *Pseudomonas syringae*, including HopZ1a, and *Xanthomonas* species. Receptor-like cytoplasmic kinases (RLCKs) such as ZED1, interact with ZAR1 and provide specificity for different effector proteins, such as HopZ1a. We previously developed a transient expression system in *Nicotiana benthamiana*, that allowed us to demonstrate ZAR1 function is conserved from the Brassicaceae to the Solanaceae. Here, we combined structural modeling of ZAR1, with molecular and functional assays in our transient system, to show that multiple intramolecular and intermolecular interactions regulate ZAR1 activity. We identified new determinants required for the formation of the ZAR^CC^ dimer and its activity. Lastly, we characterized new intramolecular interactions between ZAR1 subdomains that participate in keeping ZAR1 immune complexes inactive. This work identifies molecular constraints on immune receptor function and activation.

**One sentence-summary:** Structure-informed analyses reveal multiple finely-tuned intramolecular interactions that regulate the activity of the immune receptor ZAR1.

**Funding:** Research on plant immunity in the Lewis laboratory was supported by the USDA ARS 2030-21000-046-00D and 2030-21000-050-00D (JDL), and the NSF Directorate for Biological Sciences IOS-1557661 (JDL). ECM and AJP acknowledge financial support from UEFISCDI grant PN-III-ID-PCE-2016-0650 and the Romanian Academy programs 1 & 2 of IBAR.

## INTRODUCTION

The plant immune system activates rapid and highly effective defense mechanisms upon detection of pathogens in the vicinity of host cells (Jones and Dangl, 2006; Wu et al., 2018). Plants detect highly conserved pathogen-associated molecular patterns (PAMPs) using cell surface-localized pattern recognition receptors (PRRs), in a process called Pattern Triggered Immunity (PTI) (Segonzac and Zipfel, 2011; Monaghan and Zipfel, 2012). Many phytopathogenic bacteria encode a molecular syringe called the type III secretion system, which injects effector proteins into the plant cell (Galán and Wolf-Watz, 2006). Effector proteins primarily function to suppress PTI and promote bacterial virulence (Mudgett, 2005; Grant et al., 2006; Block et al., 2008; Lewis et al., 2009; Deslandes and Rivas, 2012; Xin and He, 2013; Macho, 2016). Plants sense pathogen activities through the cytoplasmic detection of effector proteins by a family of proteins that contain a nucleotide-binding-Apaf1-R protein-CED4 (NBARC) domain and a leucine-rich repeat domain (LRR) and are termed NLR proteins (Schreiber et al., 2016a; Khan et al., 2017; Urbach and Ausubel, 2017; Jones et al., 2016). Effector-triggered immunity (ETI) is particularly strong and often results in a form of localized programmed cell death called the hypersensitive response (HR) (Heath, 2000).

Plants encode numerous NLR genes, which use different well-characterized strategies to protect plants against pathogens (Kourelis and van der Hoorn, 2018) (Cesari, 2018). The most straightforward way of detecting the effector is the physical binding of the NLR protein with the microbial effector following a ligand-receptor model (Cesari, 2018). However, in most cases, the recognition is indirect and involves another host protein called the “guardee” that is monitored by the NLR protein. The “guardee” can be a virulence target that is modified by the effector to promote disease in the absence of the NLR protein (the guard model) (Khan et al., 2016; Dangl and Jones, 2001; Van Der Biezen and Jones, 1998; Schreiber et al., 2016a). In some cases, the plant evolves a protein that mimics a virulence target (a “decoy”) to trap the effector; here the decoy is not targeted by the pathogen to promote host susceptibility in the absence of the NLR protein (decoy model) (van der Hoorn and Kamoun, 2008; Cesari, 2018; Schreiber et al., 2016a). In addition, some NLR proteins encode an additional domain that mimics the effector target (integrated domain/decoy model) (Kroj et al., 2016; Baggs et al., 2017).

In addition to the NBARC and LRR domains, NLRs typically contain an additional N-terminal domain, a Toll/interleukin1 receptor (TIR) or coiled coil (CC) domain, which roughly define two main groups of NLRs (McHale et al., 2006; Meyers et al., 2003). These N-terminal domains are often associated with triggering downstream signaling as the overexpression of several CC and TIR domains alone induces a HR (Frost et al., 2004; Swiderski et al., 2009; Williams et al., 2014; Wróblewski et al., 2018). HR activation by CC or TIR domains has been correlated, in some cases, with homodimerization, which has also been observed in mammalian NLRs (Maekawa et al., 2011; Bernoux et al., 2011; Bentham et al., 2017; Zhang et al., 2017; Jones et al., 2016). The NBARC domain acts as a molecular switch, binding ADP in the inactive state and ATP in the active state of the protein (Wendler et al., 2012; Sukarta et al., 2016). Mutations that affect the nucleotide-bound state of the NBARC domain often result in auto-activity or loss of function, which demonstrates the crucial role of this domain (Tameling et al., 2006; Tameling et al., 2002; Bendahmane et al., 2002; DeYoung and Innes, 2006). The C-terminal LRR domain interacts with the effector in the case of direct recognition (Schreiber et al., 2016b; Ravensdale et al., 2012), or with the “guardee” in cases of indirect recognition (Baudin et al., 2017; Qi et al., 2012; Collier and Moffett, 2009). Since NLR-activated downstream signaling often results in cell death and a high fitness cost for the plant, NLRs need to be tightly regulated in the absence of pathogens. NLRs are maintained in an “off” state by intra-molecular interactions (Takken and Goverse, 2012), including between the ARC2 subdomain of the NBARC and the N-terminal part of the LRR (Slootweg et al., 2013; Sukarta et al., 2016; Rairdan et al., 2008; Ade et al., 2007; Leister et al., 2005), and between the CC or TIR domains and the NBARC domain (Wang et al., 2015a; Qi et al., 2012; Moffett et al., 2002). Structure-function analyses of several receptors led to the development of models for NLR activation based on the equilibrium between the ADP and ATP bound states (Bernoux et al., 2016; Takken and Goverse, 2012; Sukarta et al., 2016; Zhang et al., 2017). Structural modeling based on the crystal structures of CC, NBS and LRR domain homologues helps provide insights into the conformational changes that occur during NLR activation (Takken and Goverse, 2012; Slootweg et al., 2013).

HOPZ-ACTIVATED RESISTANCE 1 (ZAR1) is a canonical CC-type NLR that was first identified in *Arabidopsis thaliana*, and is involved in the immune responses against HopZ1a and HopF2a, effectors from the phytopathogenic bacteria *Pseudomonas syringae* (Lewis et al., 2010; Seto et al., 2017) and AvrAC, an effector from the phytopathogenic bacteria *Xanthomonas campestris* (Wang et al., 2015b). Recognition of diverse effectors is possible due to the association of ZAR1 with Receptor-Like Cytoplasmic Kinases (RLCK) that are encoded in a genomic cluster in Arabidopsis and that serve as adaptors/baits for the effectors (Lewis et al., 2013; Wang et al., 2015b; Seto et al., 2017). HopZ1a triggers a strong HR, that depends on the RLCK ZED1 (HOPZ-EFFECTOR-TRIGGERED IMMUNITY DEFICIENT 1) and ZAR1 (Lewis et al., 2008, 2010, 2013; Baudin et al., 2017). HopZ1a acetylates ZED1 which is hypothesized to trigger ZAR1 activation, likely through conformational changes (Lewis et al., 2013; Baudin et al., 2017). The recognition of AvrAC and HopF2a is more complex and involves other proteins that mediate a connection between the effector and the ZAR1-RLCK complex (Wang et al., 2015b; Seto et al., 2017). In the case of AvrAC, the effector interacts with and uridylylates the PBL2 (PBS1-LIKE 2) kinase which then interacts with the RLCK RKS1 (RESISTANCE RELATED KINASE 1) and ZAR1 (Wang et al., 2015b). For HopF2a, the ZRK3 (ZED1-RELATED KINASE 3) RLCK does not interact directly with the effector, which suggests that another RLCK like PBL2 might be involved (Seto et al., 2017). *N. benthamiana* ZAR1 (NbZAR1) was recently identified together with the RLCK JIM2 (XOPJ4 IMMUNITY 2) as being required for the recognition of the *Xanthomonas perforans* effector XopJ4, which is a member of the HopZ/YopJ superfamily of effector proteins (Lewis et al., 2009; Schultink et al., 2019). Spurious activation of ZAR1 is, in part, controlled by intermolecular interactions with ZED1/RKS1/ZRK3/JIM2 and intramolecular interactions within ZAR1 (Baudin et al., 2017). The downstream signaling activated by ZAR1 is still uncharacterized, and acts independently of most pathways identified so far (Lewis et al., 2010; Macho et al., 2010).

In this paper, we used structural modeling together with molecular assays and functional studies to decipher the interactions and conformational changes that regulate ZAR1 activity. We previously established a *N. benthamiana* transient expression system that allowed us to evaluate the contributions of ZAR1 and ZED1 to HopZ1a recognition and demonstrated that ZAR1-mediated recognition is conserved from the Brassicaceae to the Solanaceae (Baudin et al., 2017). We took advantage of this system to evaluate the impact of domain truncations and loss-of-function mutations on ZAR1 intramolecular interactions and activity. Finally, our interdomain contact data provide new insights into our understanding of ZAR1 architecture. This work will help identify the constraints on immune receptor function and activation.

## RESULTS

### Model-based identification of ZAR1^CC^ structural determinants for homodimerization

We previously showed that the overexpression of the CC domain of ZAR1 (ZAR1^CC^ amino acids 1-144) induces a HR in *N. benthamiana* when ZAR1^CC^ was cloned as an in-frame fusion to YFP (Baudin et al., 2017). We further demonstrated that ZAR1^CC^ and a shorter version of ZAR1^CC^ (ZAR1^CC2^, amino acids 1-121) homodimerize in planta (ZAR1^CC^) and in yeast (ZAR1^CC^ or ZAR1^CC2^) (Baudin et al., 2017). The CC domains of several NLRs, including MLA10 (MILDEW LOCUS A 10) from *Hordeum vulgare* (barley) and Rp1 (RESISTANCE to PUCCINIA 1) from *Zea mays* (maize), also homodimerize and are autoactive when expressed alone or as a fusion with YFP (Maekawa et al., 2011; Wang et al., 2015a; Bai et al., 2012).

To better understand the structural dynamics that lead to ZAR1^CC^ dimerization and autoactivation, we examined the effects of mutations introduced into critical parts of the CC structure identified by molecular modeling. Based on Rx [RESISTANCE to PVX, *Solanum tuberosum* (potato)] and Sr33 [STEM RUST 33, *Triticum aestivum* (wheat)] templates, ZAR1^CC^ can be modeled in the monomeric state as a four short helices bundle structure interconnected by three turns (1CC4α: α1a-t1-α1b-t2-α2-t3-α3) (Figure 1A, S1A, S1B). Based on the MLA10 template, the dimeric state can be modeled as two intertwined long helices, where the second helix is broken by a short coil break, separated by only one central turn (2CC2α: α1-t2-α2-c-α3) (Figure 1A, S1B). The two peripheral short helices of 1CC4α (α1a and α3) perfectly superimpose the beginning of α1 and α3 in the second monomer of the dimer (Casey et al., 2016) (Figure 1A). This suggests that the regions corresponding to t1 and t3 in the 1CC4α structure might play a role in dimerization. In addition, the first helix in the 1CC4α monomeric state is the only helix region of the CC domain that is not constrained at both ends by the rest of ZAR1, and is free to move at its N-terminal end. Therefore, its hydrophobic zipping to the rest of the CC domain, and the propensity of α1a and α1b to form a turn or a helix might be critical in dimerization or protein-protein interactions.

**Figure 1.**
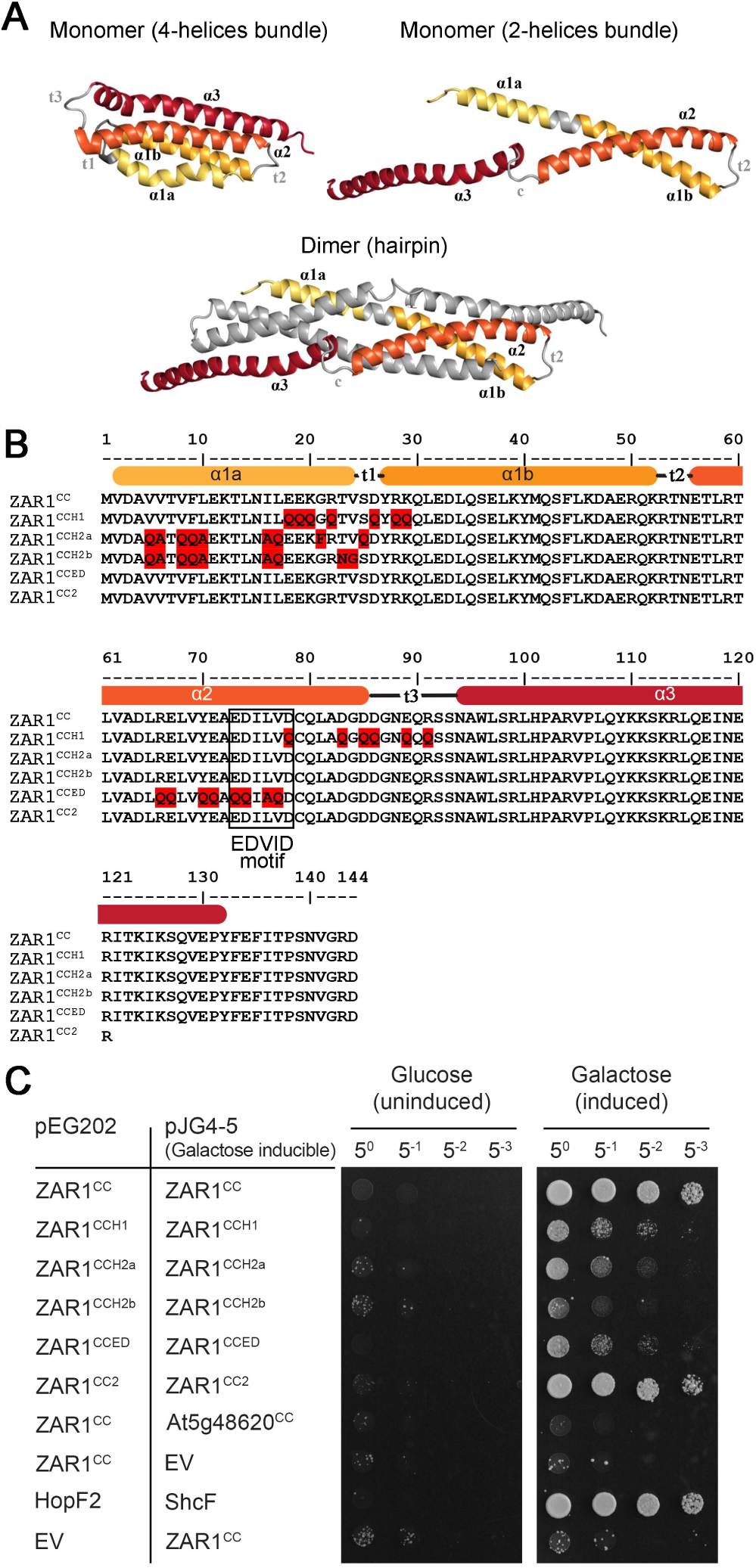
Effect of motifs in ZAR1^CC^ on dimerization. A, Predicted structural representation of a four-helices bundle monomer, a two-helices bundle monomer and a hairpin dimer. The different helical segments are indicated as α1a, α1b, α2 and α3. The 4 consecutive helices are colored from bright orange to dark red, and the second molecule in the dimer is shown in grey. B, Sequence alignment of ZAR1^CC^ and mutants in the CC domain are shown with a schematic representation of the helical domains. Colored residues were mutated to the residues shown in each construct. The EDVID motif is boxed. C, Yeast two-hybrid analysis of ZAR1^CC^ dimerization. Sequences in the pEG202 vector are constitutively expressed as fusions to a LexA DNA-binding domain, while pJG4-5 confers galactose-inducible expression of sequences as fusions to a B42 transcriptional activation domain. The CC domain from an unrelated Arabidopsis NLR (At5g48620) was used as a negative control, while the genes HopF2 and ShcF from *Pseudomonas syringae* pv. tomato DC3000 are a positive control, having been previously shown to strongly interact (Shan et al., 2004).

To better understand the role of different motifs in the CC domain, we generated five versions of ZAR1^CC^ (hereafter ZAR1^CCH1^, ZAR1^CCH2a^, ZAR1^CCH2b^, ZAR1^CC2^, ZAR1^CCED^) (Figure 1B, S1C). ZAR1^CCH1^ has mutations in the lateral loops, in which the amino acids that are predicted to form salt bridges have been mutated. To reduce the probability of transient interactions between the two monomers at the α1a-α1b/α2-α3 site, we introduced glutamine mutations which eliminated the favorable charge-charge interaction network between two monomers in that region. For ZAR1^CCH2a^ and ZAR1^CCH2b^, hydrophobic amino acids of the first helix (α1a) were replaced to investigate whether increased mobility at α1a-α1b region affects the mechanism of dimerization. For ZAR1^CCH2a^, the first loop (t1) was modified to increase the local helix propensity and stabilize the interaction. As in the MLA10 dimeric structure, this region corresponds to an uninterrupted α1a-α1b helix. For ZAR1^CCH2b^, the loop region between t1a and t1b was modified to be less prone in adopting a helical fold and therefore should be destabilizing. ZAR1^CCED^ targets the conserved EDVID motif (amino acids 73-77), which is found in many CC-type NLRs (Bai et al., 2012; Rairdan et al., 2008; Baudin et al., 2017; El Kasmi et al., 2017) (Figure S1A, S1C). Finally, we generated a shorter version of the CC domain, ZAR^CC2^ that can still dimerize in yeast (Baudin et al., 2017).

To independently test the dimerization capacity of each construct, we conducted yeast two-hybrid assays (Y2H). We cloned the five versions of ZAR1^CC^ as a fusion to the activation domain or DNA-binding domain in the LexA yeast two-hybrid system. For each combination, the same optical density of yeast was spotted on the plates to quantitate the interaction strength. The ZAR1^CCH1^, ZAR1^CCH2a^, and ZAR1^CCED^ mutations showed similar levels of yeast growth that were weaker than that of wild type ZAR1^CC^ (Figure 1C). The ZAR1^CCH2b^ mutations had the most substantial effect on yeast growth compared to the other constructs (Figure 1C). Mutation of the EDVID motif only slightly reduced the interaction strength, indicating that this motif is not absolutely required for dimerization (Figure 1C). ZAR1^CC^ did not interact with an unrelated NLR, At5g48620, indicating that homodimerization is specific (Figure 1C). The lack of interactions was not due to a lack of expression (Figure S2).

Our data indicate that CC dimerization is impacted by mutations which affect charge-charge interactions around t1 and t3 in ZAR1^CCH1^, or by disrupting the hydrophobic zipping in α1a in ZAR1^CCH2a^ and ZAR1^CCH2b^. The mutations in ZAR1^CCH2b^ that increase local flexibility of the α1a-α1b region have a stronger effect on shifting the equilibrium towards the monomeric state, compared to the mutations in ZAR1^CCH2a^ that increase the helical propensity within the same region.

### Impact of ZAR1^CC^ architecture on its activity

The ability of ZAR1^CC^ to induce autoactivity is dependent on the presence of a YFP epitope, suggesting that the weak dimerization capacity of YFP stabilizes ZAR1^CC^ oligomerization (Baudin et al., 2017). However, no clear correlation between ZAR1^CC^ dimerization and autoactivity has been demonstrated so far. We took advantage of the mutations we generated in the ZAR1^CC^ domain (Figure 1B) to test their capacity for autoactivity as a ZAR1^CC^-YFP fusion. When over-expressed in *N. benthamiana* leaves, the YFP fusions of ZAR1^CCH1^, ZAR1^CCH2a^, ZAR1^CCH2b^, ZAR1^CC2^ and ZAR1^CCED^ did not lead to autoactivity, while wild type ZAR1^CC^-YFP still induced autoactivity (Figure 2A). The absence of a HR-like phenotype was not due to the absence of expression as all constructs are equally expressed (Figure 2B). Consistent with reduced dimerization in ZAR1^CCH1^ or ZAR1^CCH2a^, or even less dimerization in ZAR1^CCH2b^, mutations in all of these constructs disrupted autoactivity of the CC domain. Interestingly, both ZAR1^CC2^-YFP and ZAR1^CCED^-YFP were also unable to induce autoactivity (Figure 2A), even though ZAR1^CC2^ could dimerize like wild type and ZAR1^CCED^ dimerized poorly (Figure 1C). These data indicate that the C-terminal portion of the CC domain and the EDVID motif are required to activate downstream signaling, and that dimerization is not sufficient to activate downstream signaling.

**Figure 2.**
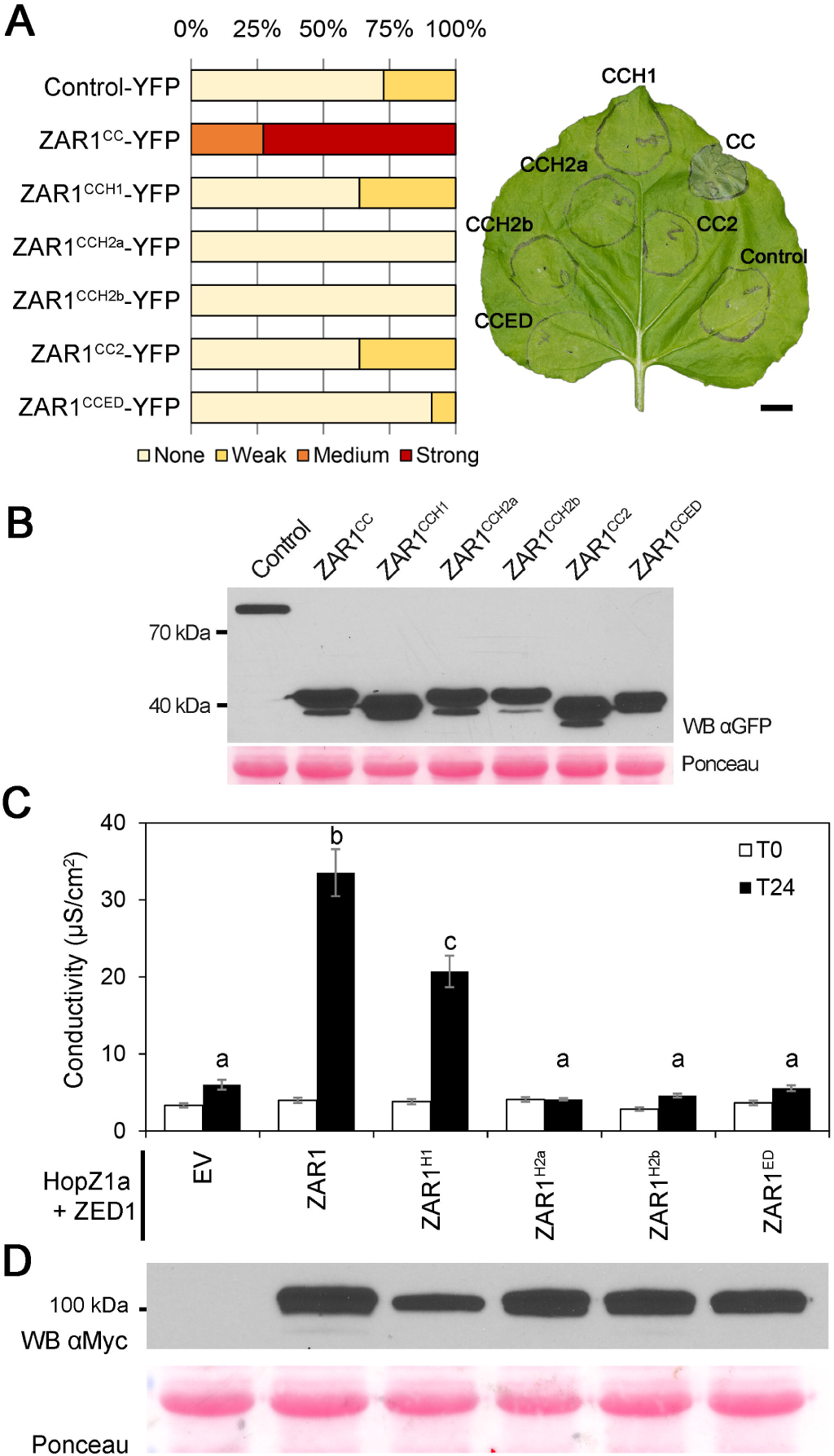
ZAR1^CC^ motifs contribute to HR when inducibly expressed in *N. benthamiana*. *Agrobacterium tumefaciens* carrying the indicated constructs were pressure-infiltrated into *N. benthamiana* leaves. Constructs were expressed under a dexamethasone-inducible promoter. A, CC domain mutants do not trigger a HR-like phenotype. ZAR1^CC^, ZAR1^CC^ mutants or a control protein (At3g46600) were expressed as fusions to YFP. Phenotypes were observed and photographs were taken 48 h after dexamethasone induction. The percentage of leaves showing the HR phenotype is shown after being scored on a scale from no HR, weak HR, medium HR to strong HR. A representative leaf with the HR phenotype is shown. The scale bar is 1 cm. The experiment was repeated 3 times with similar results. B, Immunoblot analysis of ZAR1^CC^-YFP protein expression. Samples were harvested 24 h after dexamethasone induction. Equal amounts of proteins were resolved on 10% SDS-PAGE gels, blotted onto nitrocellulose, and probed with GFP antibodies in Western blot analysis (WB). The Ponceau red stained blot was used as the loading control. C, Full-length ZAR1 with mutations in the CC domain exhibit partial or no complementation of *NbZAR1*-VIGS. Levels of ion leakage induced by ZAR1, ZAR1 mutants, or EV were measured at 4 h (T0) and 28 h (T24) after dexamethasone application. Error bars indicate the standard error from the mean of 6 samples. The letters beside the bars indicate significance groups, as determined by a one-way ANOVA comparison followed by a Tukey’s post hoc test (*P*value ≤ 0.05). The experiment was repeated 3 times with similar results. D, Immunoblot analysis of ZAR1 protein expression. Samples were harvested 24 h after dexamethasone induction. Equal amounts of proteins were resolved on 10% SDS-PAGE gels, blotted onto nitrocellulose, and probed with Myc antibodies in Western blot analysis (WB). The Ponceau red stained blot was used as the loading control.

To analyze the effect of these ZAR1^CC^ mutations on ZAR1 function, we introduced the ZAR1^CC^ mutations into full-length ZAR1 (hereafter ZAR1^H1^, ZAR1^H2a^, ZAR1^H2b^, and ZAR1^ED^) and tested them for complementation of *NbZAR1* silencing in *N. benthamiana* as previously described (Baudin et al., 2017). HopZ1a and ZED1 were co-expressed with the ZAR1 mutants, ZAR1 as a positive control, or empty vector as a negative control, in *N. benthamiana* leaves silenced for *NbZAR1* by virus-induced gene silencing (VIGS). To quantify the HR intensity, and thus the complementation capacity of each construct, we measured ion leakage into the media over time. Wild type ZAR1 showed high levels of conductivity (Figure 2C), as previously observed (Baudin et al., 2017). ZAR1^H1^ showed moderate conductivity that was weaker than and statistically different from wild type ZAR1 (Figure 2C). The ZAR1^H2a^, ZAR1^H2b^ and ZAR1^ED^ constructs resulted in conductivity measurements that were similar to empty vector (Figure 2C). Therefore, ZAR1^H1^ was able to partially complement *NbZAR1*-VIGS while ZAR1^H2a^, ZAR1^H2b^ and ZAR1^ED^ did not display any complementation of *NbZAR1*-VIGS (Figure 2C). The differences observed in the conductivity assay were not due to differences in expression level as all the ZAR1 constructs accumulated to similar levels (Figure 2D).

To investigate if ZAR1^CC^-YFP-induced autoactivity requires the presence of the endogenous *NbZAR1*, we expressed ZAR1^CC^-YFP in silenced *NbZAR1-VIGS* or control *GUS*-VIGS plants. We observed a strong HR in both *GUS*-VIGS and *NbZAR1*-VIGS plants indicating that endogenous *NbZAR1* is not needed for ZAR1^CC^-YFP autoactivity (Figure S3). VIGS silencing was efficient because co-expressed HopZ1a and ZED1 were not able to induce the HR in *NbZAR1*-VIGS plants while they showed a strong HR in *GUS*-VIGS plants (Figure S3).

Taken together, these data indicate that the first helix (α1a) and the EDVID motif of ZAR1^CC^ are particularly important as mutations in these regions lead to a non-functional ZAR1 protein. The mutations in the ZAR1^H1^ construct have a less significant impact, as ZAR1^CCH1^ was still able to dimerize and ZAR1^H1^ was still able to partially complement *NbZAR1*-VIGS. In addition, ZAR1^CC^-YFP autoactivity is independent of *NbZAR1*.

### Autoactivity of ZAR1^CC^-YFP is inhibited by the ZAR1^NBARC^ domain

We previously showed that ZAR1^CCNBARC1^-YFP did not elicit a HR-like phenotype, indicating that the NBARC1 domain (amino acids 145-391) suppresses autoactivity of the CC domain (Baudin et al., 2017). We hypothesized that the inhibitory effect of ZAR1^NBARC^ on ZAR1^CC^ autoactivity was due to an intramolecular interaction between these two domains. To test this hypothesis, we performed Y2H assays between ZAR1^CC^ and different ZAR1^NBARC^ truncations. We constructed ZAR1^NBARC^ variants that contained the NB subdomain alone (ZAR1^NB^, amino acids 145-315), the NB subdomain and the first ARC subdomain (ZAR1^NBARC1^, amino acids 145-391) or the NB subdomain and both ARC subdomains (ZAR1^NBARC1+2^, amino acids 145-504), as in-frame fusions to the activation domain. Each was tested for an interaction with ZAR1^CC^ cloned as an in-frame fusion to the DNA-binding domain in yeast. We observed significant yeast growth for all three truncations of the NBARC domain when co-expressed with ZAR1^CC^ but not with the negative control At5g48620^CC^ (Figure 3A). The full-length NBARC domain (ZAR1^NBARC1+2^) showed the strongest interaction with ZAR1^CC^ (Figure 3A). All constructs were expressed at similar levels in yeast (Figure S2).

**Figure 3.**
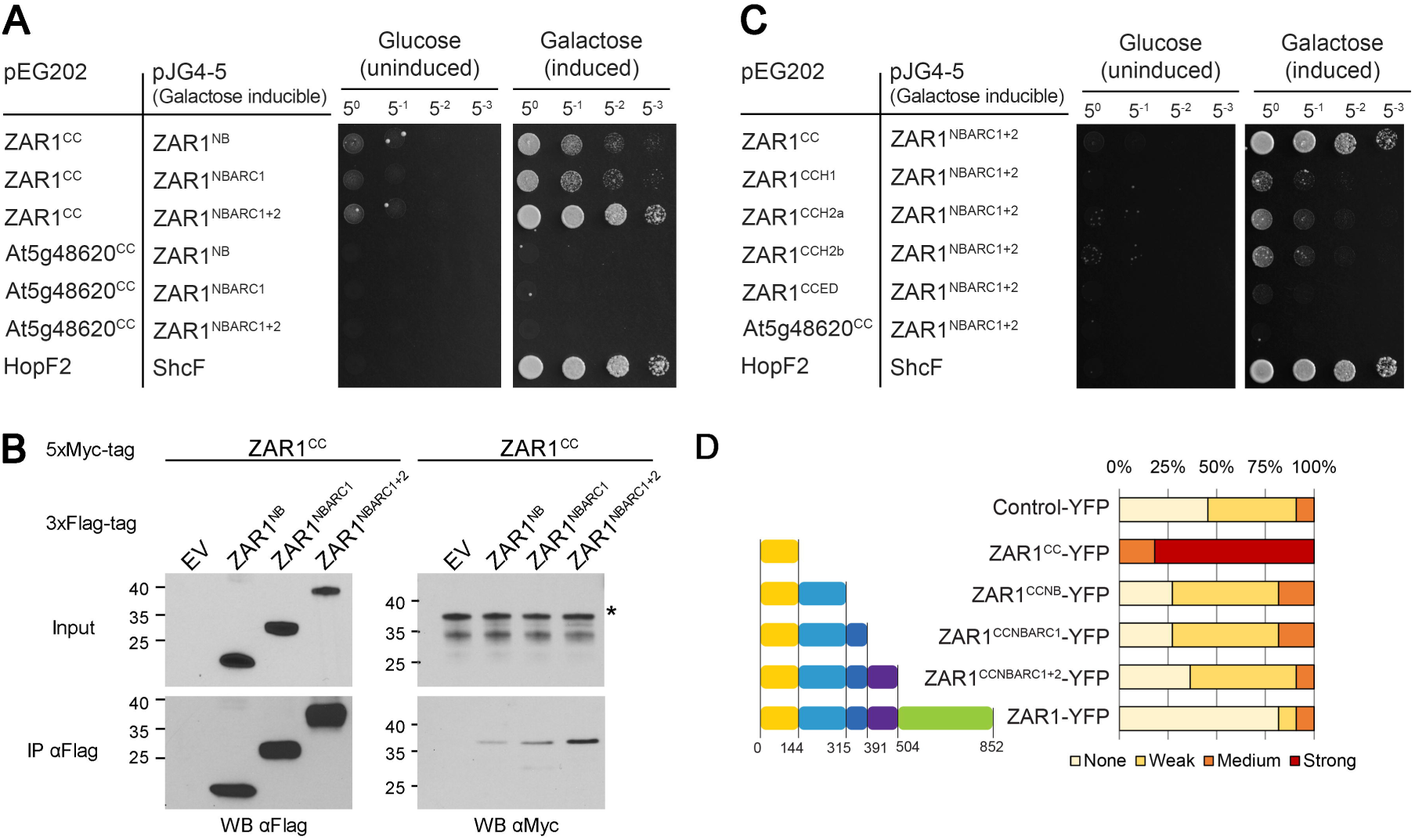
The CC and NBARC domains interact and the NBARC domain suppresses autoactivity. A, Yeast two-hybrid analysis of ZAR1^CC^ interactions with varying regions of the NBARC domain. NB indicates that only the NB subdomain is present. NBARC1 indicates that the NB and ARC1 subdomains are present. NBARC1+2 indicates that the NB, ARC1 and ARC2 subdomains are present. Sequences in the pEG202 vector are constitutively expressed as fusions to a LexA DNA-binding domain, while pJG4-5 confers galactose-inducible expression of sequences as fusions to a B42 transcriptional activation domain. The CC domain from an unrelated Arabidopsis NLR (At5g48620) was used as a negative control, while the genes HopF2 and ShcF from *Pseudomonas syringae* pv. tomato DC3000 provided a positive control, having been previously shown to strongly interact (Shan et al., 2004). B, Immunoblot showing the coimmunopurification of ZAR1^CC^ with ZAR1^NB^, ZAR1^NBARC1^ and ZAR1^NBARC1+2^ from *N. benthamiana* tissues. ZAR1^CC^ with a 5xMyc epitope tag was coexpressed with empty vector (EV) or NBARC variants with a 3xFlag epitope tag in *N. benthamiana*. Western-blot analysis (WB) was performed on the crude extract (input) or the immunopurified fractions (IP) from anti Flag beads. An asterisk indicates the band corresponding to the protein of interest, if multiple bands were observed. This experiment was repeated three times with similar results. C, Yeast two-hybrid analysis of interactions between ZAR1^CC^ mutants and ZAR1^NBARC1+2^, as in A. D, The NB domain of ZAR1 is sufficient to suppress autoactivity of the CC domain. ZAR1^CC^, ZAR1^CCNBARC^ truncations, ZAR1 or a control protein (At3g46600) were expressed as fusions to YFP. The colored boxes correspond to the following domains: yellow is the CC domain, medium blue is the nucleotide-binding (NB) subdomain, dark blue is the Apaf1-R protein-CED4 1 (ARC1) subdomain, purple is the ARC2 subdomain, and green is the leucine-rich repeat (LRR) domain. The percentage of leaves showing the HR phenotype is shown after being scored on a scale from no HR, weak HR, medium HR to strong HR. Phenotypes were observed 48 h after dexamethasone induction. The experiment was repeated 3 times with similar results.

To confirm these interactions *in planta*, we carried out co-immunoprecipitation experiments (CoIP). We cloned NBARC variants as in-frame fusions with a 3xFlag epitope and ZAR1^CC^ as an in-frame with a 5xMyc epitope. We co-expressed ZAR1^CC^-5xMyc with each NBARC variant or empty vector (EV) in *N. benthamiana* leaves, followed by immunoprecipitation of protein complexes using anti-Flag magnetic beads. As seen in yeast (Figure 3A), ZAR1^CC^ interacted with all three variants and showed the strongest interaction with ZAR1^NBARC1+2^ (Figure 3B). Lastly, we confirmed these interactions using bimolecular fluorescence complementation (BiFC) in *N. benthamiana*. We cloned NBARC variants as in-frame fusions with the N-terminal part of YFP (n-YFP) and ZAR1^CC^ as an in–frame fusion with the C-terminal part of YFP (cYFP), and co-expressed these constructs in *N. benthamiana* leaves. We observed strong fluorescence when ZAR1^CC^-cYFP was co-expressed with all three variants of the NBARC domain which further validates the Y2H and CoIP experiments and confirms that the CC and NBARC domains of ZAR1 interact (Figure S4A).

To explore structural determinants in the CC domain that might be required for the interaction with ZAR1^NBARC^, we then tested for an interaction between our ZAR1^CC^ mutants (Figure 1B) and ZAR1^NBARC1+2^ in Y2H assays. While wild type ZAR1^CC^ was able to interact with ZAR1^NBARC1+2^ as we showed previously, ZAR1^CCH1^, ZAR1^CCH2a^ and ZAR1^CCH2b^ were strongly impaired in their ability to interact with ZAR1^NBARC1+2^ (Figure 3C). Interestingly, ZAR1^CCED^ was completely unable to interact with ZAR1^NBARC1+2^, suggesting that this motif plays a particularly important role in the intramolecular interaction.

To further delineate what part of the NBARC domain was necessary for suppression of ZAR1^CC^ autoactivity, we cloned C-terminal truncations of ZAR1 as in-frame fusions with YFP and expressed them under a dexamethasone-inducible promoter in *N. benthamiana*. We compared ZAR1^CC^ with ZAR1^CCNBARC^ variants that contained the CC and NB domains (ZAR1^CCNB^, amino acids 1-315), the CC, NB and first ARC subdomain (ZAR1^CCNBARC1^, amino acids 1-391) or the CC, NB and both ARC domains (ZAR1^CCNBARC1+2^, amino acids 1-504) (Figure 3C). ZAR1^CC^-YFP induced a strong HR-like phenotype, while ZAR1^CCNB^-YFP, ZAR1^CCNBARC1^-YFP and ZAR1^CCNBARC1+2^-YFP did not display a HR-like phenotype (Figure 3D). These data indicate that the NB subdomain is sufficient to suppress ZAR1^CC^ autoactivity. The absence of autoactivity was not due to the absence of protein expression as all the constructs showed similar expression levels (Figure S4B).

Taken together, our data show that the ZAR1^CC^ and ZAR1^NBARC1+2^ interaction requires the EDVID motif in ZAR1^CC^, and that proper folding of the ZAR1^CC^ domain contributes to interactions with ZAR1^NBARC1+2^. As well, the NB domain alone was sufficient to suppress the autoactivity of ZAR1^CC^-YFP.

### Effect of loss- and gain-of-function mutations on the ZAR1^NBARC^ domain

We and others previously identified loss-of-function mutations in *zar1* that resulted in a lack of HopZ1a or AvrAC recognition (Lewis et al., 2013; Wang et al., 2015b; Baudin et al., 2017). Among these loss-of function mutations, K195N, V202M and S291N are located in the NB subdomain, P359L is in the ARC1 subdomain, and L465F is in the ARC2 subdomain (Figure 4A). To determine if these mutations in the NBARC domain affected the interaction with ZAR1^CC^, we constructed ZAR1^NBARC1+2^ variants carrying each EMS mutation with an in-frame 3xFlag tag, and co-expressed them with ZAR1^CC^-5xMyc in CoIP assays. We found that ZAR1^CC^ coimmunoprecipitated with all of them, indicating that these substitutions did not affect the interaction between the CC and NBARC domain of ZAR1 (Figure 4B). Since the CC domain could interact with the NBARC domain carrying each mutation, we tested whether the mutated NBARC domain could still suppress autoactivity of the CC domain. To maximize any effect of the mutation, we used the minimally interacting NBARC domain that would still include each EMS mutation. The three NB mutations (K195N, V202M and S291N) were cloned into ZAR1^CCNB^ with an in-frame C-terminal YFP tag. The ARC1 mutation (P359L) was cloned into ZAR1^CCNBARC1^ with an in-frame C-terminal YFP tag. The ARC2 mutation (L465F) was cloned into ZAR1^CCNBARC1+2^ with an in-frame C-terminal YFP tag. None of the constructs showed autoactive phenotypes as displayed by the positive control ZAR1^CC^-YFP when overexpressed in *N. benthamiana* leaves (Figure 4C). The absence of phenotype was not due to the absence of expression as all of the constructs expressed well in *N. benthamiana* (Figure S4C). Therefore, the five loss-of function mutations in ZAR1^NBARC1+2^ do not play a role in ZAR1^CC^ intramolecular interactions or in NBARC suppression of ZAR1^CC^-YFP autoactivity.

**Figure 4.**
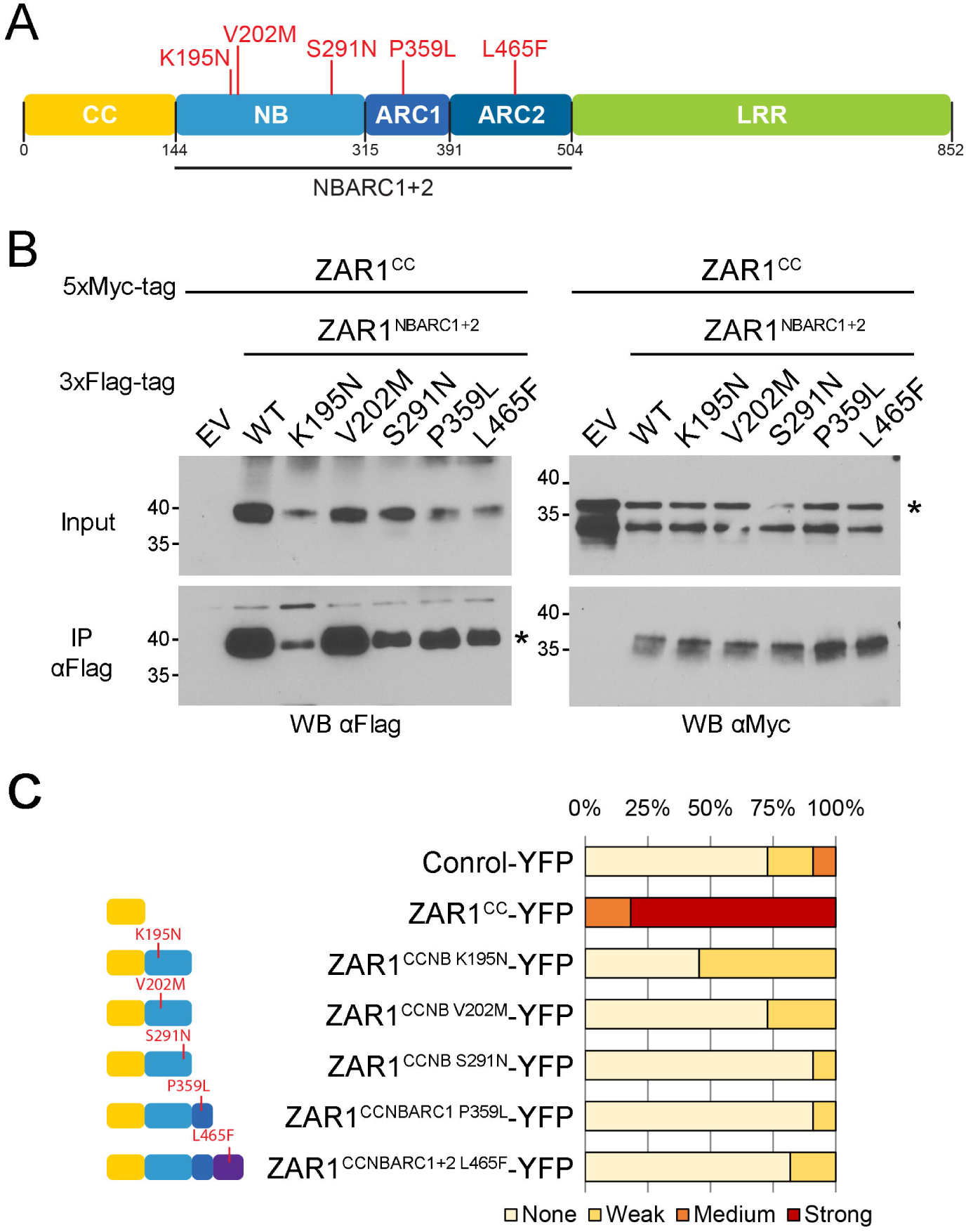
Mutations in the NBARC domain do not impair interactions with ZAR1^CC^ or ZAR1^CC^-YFP autoactivity. A, ZAR1 sequence schematic showing the three main domains. The colored boxes correspond to the following domains: yellow is the CC domain, medium blue is the nucleotide-binding (NB) subdomain, dark blue is the Apaf1-R protein-CED4 1 (ARC1) subdomain, purple is the ARC2 subdomain, and green is the leucine-rich repeat (LRR) domain. The amino acid changes induced by the mutation are indicated above the schematic. B, Immunoblot showing the coimmunopurification of ZAR1^CC^ with ZAR1^NBARC1+2^ wild type (WT) or carrying mutations from *N. benthamiana* tissues. ZAR1^CC^ with a 5xMyc epitope tag was coexpressed with empty vector (EV) or NBARC variants with a 3xFlag epitope tag in *N. benthamiana*. Western-blot analysis (WB) was performed on the crude extract (input) or the immunopurified fractions (IP) from anti Flag beads. An asterisk indicates the band corresponding to the protein of interest, if multiple bands were observed. This experiment was repeated three times with similar results. C, ZAR1^CC^, ZAR1^CCNBARC^ truncations carrying mutations, or a control protein (At3g46600) were expressed as fusions to YFP. The percentage of leaves showing the HR phenotype is shown after being scored on a scale from no HR, weak HR, medium HR to strong HR. Phenotypes were observed 48 h after dexamethasone induction. The experiment was repeated twice with similar results.

In an effort to identify potential autoactive mutants of ZAR1, we mined the literature for mutations in the NBARC domain that conferred autoactivity in other NLRs. The following sites were identified as strong candidates for autoactivity: G194/K195 in the P-loop from a large-scale screen of autoactive NLRs (Lolle et al., 2017), D283 in the Walker B/kinase2 motif from the tomato NLR I-2 (DeYoung and Innes, 2006; Tameling et al., 2006), and the aspartate in the MHD motif in multiple NLRs (DeYoung and Innes, 2006; Bendahmane et al., 2002; De La Fuente Van Bentem et al., 2005; Dodds et al., 2006; Howles et al., 2005; Roberts et al., 2013; Williams et al., 2011). We mutated the corresponding residues in ZAR1: G194A/K195A in the P-loop, D268E in the Walker B/kinase2 motif, D489V in the MHD motif, and tested whether they induced a HR in *N. benthamiana*. Wild type AtZAR1 did not induce a HR-like phenotype, as previously observed (Figure S5A, S5B) (Baudin et al., 2017). AtZAR1^G194E/K195A^, AtZAR1^D268E^, and AtZAR1^D489V^ also did not cause a HR-like phenotype, and all of the proteins were expressed at similar levels (Figure S5B, S5C).

### Interaction between the NBARC and LRR domains

The NBARC-LRR interaction is also important for maintaining the NLR in the “off” state, as seen for Rx (Slootweg et al., 2013; Bendahmane et al., 2002), RPS5 (RESISTANCE TO PSEUDOMONAS SYRINGAE 5) (Ade et al., 2007; Qi et al., 2012) and RPP1 (RESISTANCE TO PERONOSPORA PARASITICA 1) (Schreiber et al., 2016b). We therefore tested if ZAR1 was similarly regulated by a NBARC-LRR interaction. We cloned ZAR1^LRR^ (amino acids 505-852) as an in–frame fusion with a 5xMyc tag and co-expressed it with different truncations of the NBARC domain as previously described. We observed the strongest interaction between ZAR1^LRR^ and ZAR1^NB^, a weaker interaction between ZAR1^LRR^ and ZAR1^NBARC1^, and the weakest interaction between ZAR1^LRR^ and ZAR1^NBARC1+2^ (Figure 5A). To confirm these interactions, we carried out BiFC assays in *N. benthamiana*. We observed a fluorescence signal when ZAR1^LRR^ was co-expressed with ZAR1^NB^, ZAR1^NBARC1^ and ZAR1^NBARC1+2^ (Figure S6).

**Figure 5.**
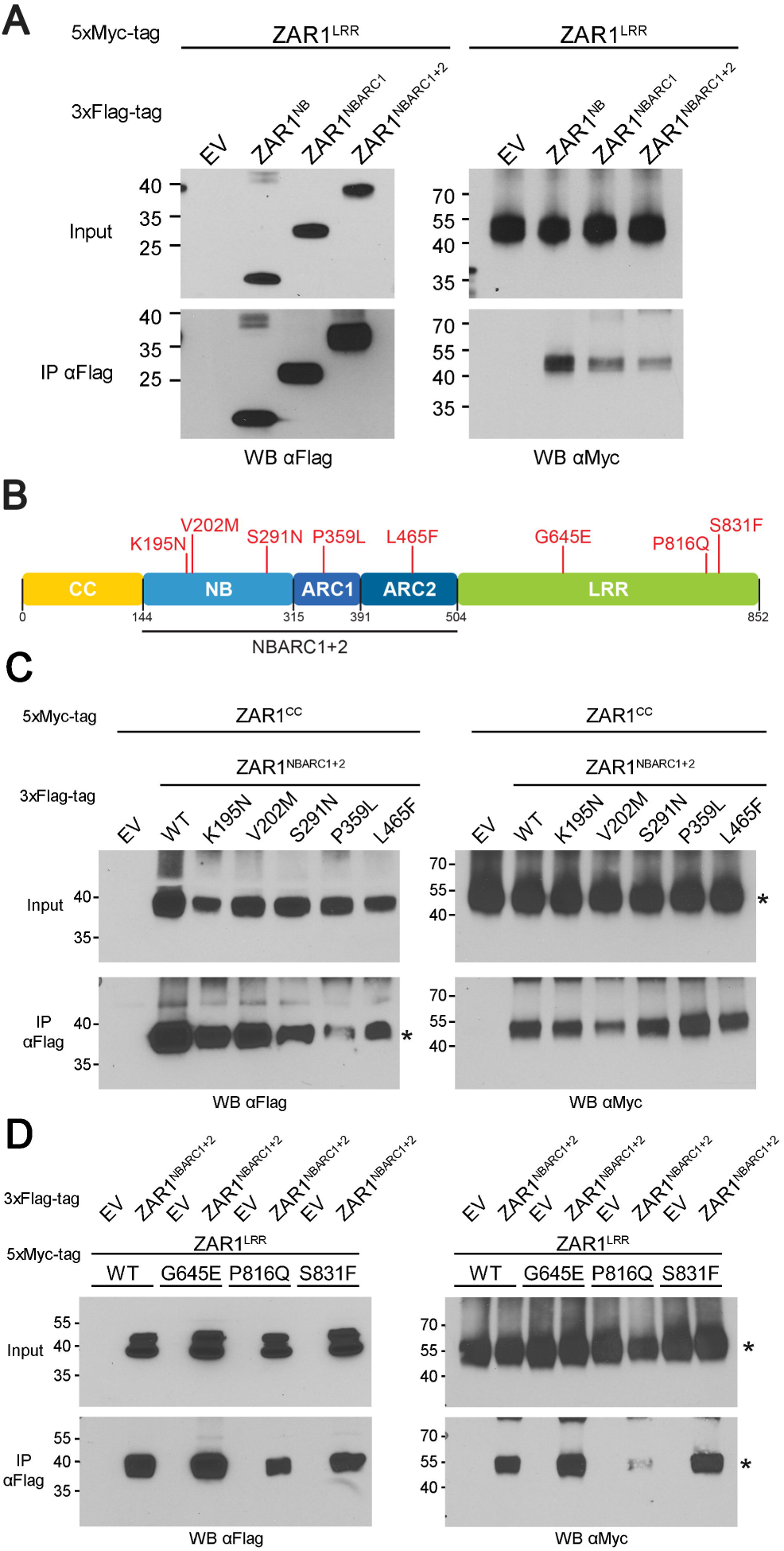
The LRR and NBARC domain of ZAR1 interact. A, Immunoblot showing the coimmunopurification of ZAR1^LRR^ with ZAR1^NB^, ZAR1^NBARC1^ and ZAR1^NBARC1+2^ from *N. benthamiana* tissues. ZAR1^LRR^ with a 5xMyc epitope tag was coexpressed with empty vector (EV) or NBARC variants with a 3xFlag epitope tag in *N. benthamiana*. NB indicates that only the NB subdomain is present. NBARC1 indicates that the NB and ARC1 subdomains are present. NBARC1+2 indicates that the NB, ARC1 and ARC2 subdomains are present. Western-blot analysis (WB) was performed on the crude extract (input) or the immunopurified fractions (IP) from anti Flag beads. This experiment was repeated three times with similar results. B, ZAR1 sequence schematic showing the three main domains. The colored boxes correspond to the following domains: yellow is the CC domain, medium blue is the nucleotide-binding (NB) subdomain, dark blue is the Apaf1-R protein-CED4 1 (ARC1) subdomain, purple is the ARC2 subdomain, and green is the leucine-rich repeat (LRR) domain. The amino acid changes induced by the mutation are indicated above the schematic. C, Immunoblot showing the coimmunopurification of ZAR1^LRR^ with ZAR1^NBARC1+2^ wild type (WT) or carrying mutations from *N. benthamiana* tissues. ZAR1^CC^ with a 5xMyc epitope tag was coexpressed with empty vector (EV) or NBARC variants with a 3xFlag epitope tag in *N. benthamiana*. Western-blot analysis (WB) was performed on the crude extract (input) or the immunopurified fractions (IP) from anti Flag beads. An asterisk indicates the band corresponding to the protein of interest, if multiple bands were observed. This experiment was repeated twice with similar results. D, Immunoblot showing the coimmunopurification of ZAR1^LRR^ wild type (WT) or carrying mutations with ZAR1^NBARC1+2^ from *N. benthamiana* tissues. ZAR1^LRR^ or mutants with a 5xMyc epitope tag were coexpressed with empty vector (EV) or ZAR1^NBARC1+2^ with a 3xFlag epitope tag in *N. benthamiana*. Western-blot analysis (WB) was performed on the crude extract (input) or the immunopurified fractions (IP) from anti Flag beads. An asterisk indicates the band corresponding to the protein of interest, if multiple bands were observed. This experiment was repeated twice with similar results.

To investigate the role of NBARC mutations on the intramolecular interaction with the LRR domain, we carried out CoIP with the ZAR1^NBARC1+2^ constructs described in Figure 4A. All of the mutants in the NBARC domain were able to coimmunoprecipitate with ZAR1^LRR^ at a similar level to wild type ZAR1^NBARC1+2^ (Figure 5C). Interestingly, the NBARC mutants were not affected in their interaction with either the CC or the LRR domains (Figure 4B, 5C), suggesting that these mutations affect distinct roles in ZAR1 function.

We then turned to mutations in the LRR domain, to evaluate their impact on the NBARC-LRR interaction. Previous genetic screens identified three loss-of-function substitutions in the LRR domain of ZAR1 (Lewis et al., 2013; Wang et al., 2015b; Baudin et al., 2017) (Figure 5B). We therefore cloned these mutations in ZAR1^LRR^-5xMyc and tested them for their ability to interact with ZAR1^NBARC1+2^-3xFlag. As we observed for the NBARC mutations (Figure 4B), none of the LRR mutations affected the interaction between ZAR1^NBARC1+2^ and ZAR1^LRR^ (Figure 5B). ZAR1^LRR-P816Q^ co-immunoprecipitated more weakly than the other LRR mutants however it was also most weakly expressed.

Taken together, these data show that the NBARC domain interacts with both the CC and LRR domains, and that the loss-of function mutations identified in the NBARC and LRR domains do not affect these intramolecular interactions.

### Remote homology modeling of ZAR1^NBARC^ and ZAR1^LRR^

To better understand potential effects of these mutations on ZAR1, we developed remote homology models of the ZAR1 NBARC domain from crystal structures of closed ADP-bound and open ATP-bound conformations of proteins in the NBARC domain superfamily. Structures included: closed and open Apaf1 from human, mouse and Drosophila (hApaf1, mApaf1 and dApaf1), open CED4 from *C. elegans*, closed and open NLRC4 from mouse, and closed NOD2 from rabbit (Riedl et al., 2005; Reubold et al., 2011; Pang et al., 2015; Cheng et al., 2016; Liu et al., 2010; Maekawa et al., 2011; Hu et al., 2013; Cheng et al., 2017; Zhang et al., 2015). Despite relatively low sequence homology (11-20% identity) among hApaf1 (5JUY - (Cheng et al., 2016)), dApaf1 (3J9L - (Pang et al., 2015)), *C. elegans* CED4 (3LQQ - (Qi et al., 2010)) and mouse NLRC4 (3JBL - (Zhang et al., 2015)), their overall 3D structure superposition was quite good. The RMSD values were ∼3Å RMSD for the hApaf1 core vs. dApaf1 and CED4, and ∼14.5Å RMSD for the hApaf1 core vs. NLRC4. When the individual NB, ARC1 or ARC2 subdomains were compared, the RMSD values were better than the aforementioned values in all cases. We used remote homology modeling to model the NBARC domain of ZAR1 based on the Apaf1 structure, as they showed the highest percentage sequence homology (∼20% identity) among all experimental NBARC structures (Figure S7, 6A). The internal structure of each subdomain was relatively similar in the ADP-bound and ATP-bound states, except for a rearrangement of the ARC2 subdomain (Figure 6A). Most loss-of-function mutations were located in or close to key motifs in the NBARC domain: K195 in the P-loop, S202 between the P-loop and RNBS-A, S291 in RNBS-B, and P359 in the GLPL motif (Figure 6A, S7).

**Figure 6.**
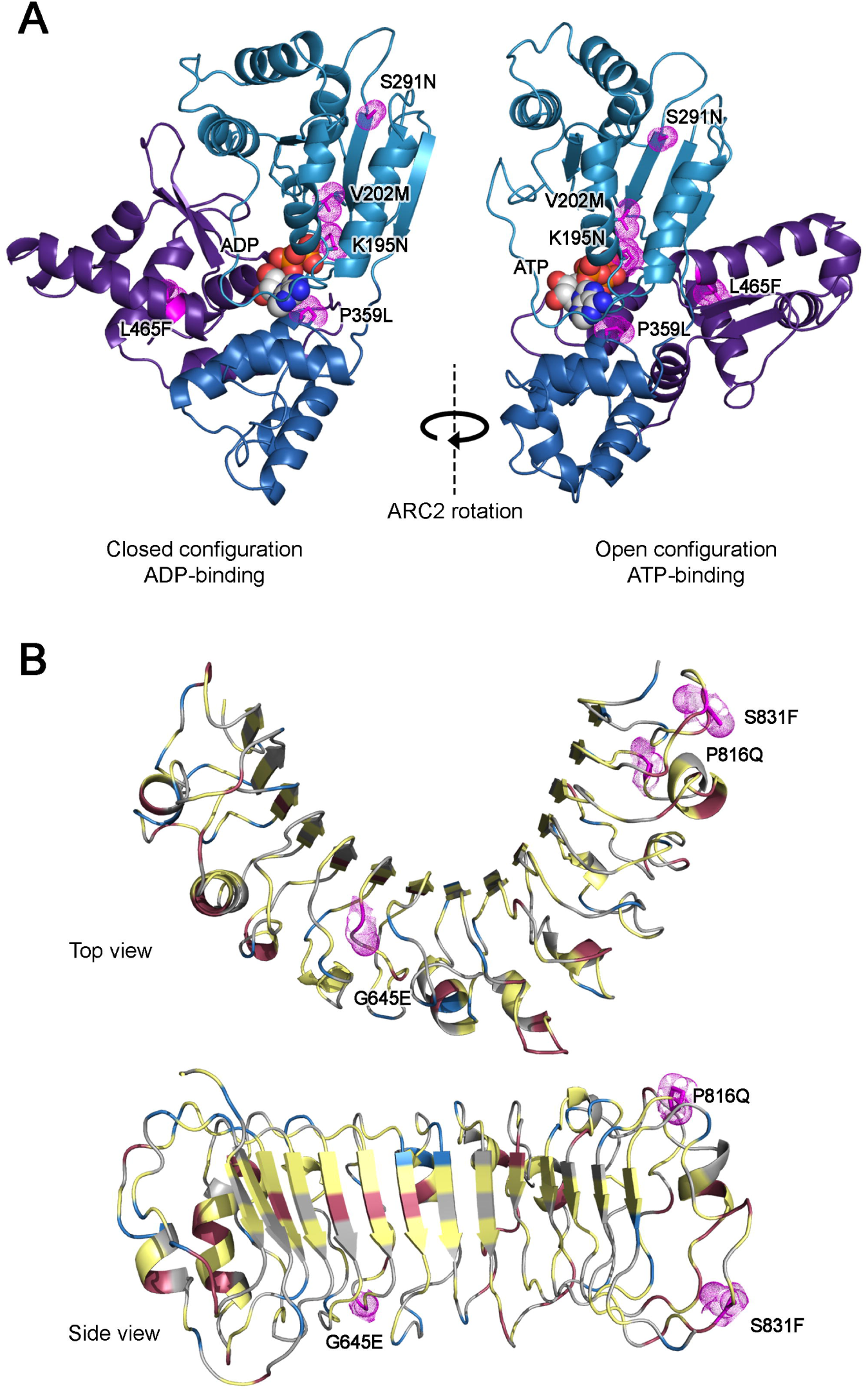
Predicted models of ZAR1^NBARC^ and ZAR1^LRR^, with mutations mapped onto the proposed 3D model. A, Models of ZAR1^NBARC^ in the closed ADP-bound conformation (left) and open ATP-bound conformation (right). NB and ARC1 were superimposed showing the rotation of ARC2 in this transition. Domains are colored as follows: medium blue is the nucleotide-binding (NB) subdomain, dark blue is the Apaf1-R protein-CED4 1 (ARC1) subdomain, and purple is the ARC2 subdomain. ADP and ATP are represented as red spheres and mutated amino acids are colored in magenta. B, The ZAR1^LRR^ domain is shown in the top view with the characteristic horseshoe shape, and as a side view for the beta sheets. Mutations are colored in magenta, hydrophobic amino acids are yellow, positively charged amino acid are blue, and negatively charged amino acids are red.

We then turned to the LRR domain of ZAR1 and used a joint fragment remote homology modeling approach, as this was shown to be effective in assessing the 3D structural properties of other LRR domains such as those in Lr10, Rx and Pm3 (Sela et al., 2012; Slootweg et al., 2013; Sela et al., 2014). Using our structural database of LRRs, the closest local ZAR1 templates were: (a) the LRR region in polygalacturonase-inhibiting protein from *Phaseolus vulgaris*, (b) the LRR protein from *Leptospira interrogans*, and (c) LRR repeats from a protein of *Agrobacterium tumefaciens* (Vorobiev et al., 2007; Di Matteo et al., 2003; Miras et al., 2015). We generated a composite model of ZAR1^LRR^ structure by joining these local fragments into the overall LRR frame based on the constraints imposed by the parallel beta strands of the ventral side of the LRR motif and a maximal structural fit over the rest of the repeat (Figure 6B, S8). The overlapping frame of the three templates runs from amino acids 510 to 823 and does not cover the last 28 residues. This analysis identified 13 LRR motifs from amino acids 510 to 823. The C-terminal region (amino acids 824-852) was modeled with low confidence based exclusively on its secondary structure propensity. Given that amino acids 843-846 are predicted in an extended configuration and are located at a distance in sequence that is compatible with one LRR repeat, the 824-852 amino acid extension was modeled here as an extra LRR repeat (Figure 6B, S8).

## DISCUSSION

To prevent cell death in the absence of pathogens, immune receptors must remain in an inactive state, while rapidly responding and transitioning to an active state upon pathogen recognition. Here, we explored the molecular determinants of ZAR1 interactions using structural modeling and functional assays in a transient assay system that we previously developed (Baudin et al., 2017). Structural modeling of ZAR1 allowed us to generate hypotheses for intramolecular interactions. We demonstrated experimentally that multiple intramolecular interactions contribute to ZAR1 function.

We carried out targeted mutagenesis of the ZAR1^CC^ domain to determine its possible structure as a monomer and how the monomer transitioned to the dimeric form. Targeted mutagenesis of ZAR1^CC^ impacted the dimerization of this sub-domain (Figures 1, S1). In particular, the ZAR1^CCH2a^ and ZAR1^CCH2b^ constructs disrupted hydrophobic zipping in α1a and shifted the monomer-dimer equilibrium in favor of the monomeric state. This is consistent with coimmunoprecipitation assays in Rx where mutations in the hydrophobic positions of the α1a region (V6E/L9E/I13E) disrupted the interaction between the 2 halves of Rx (segment α1a-α1b and segment α2-α3) (Slootweg et al., 2018). However, yeast two-hybrid assays with MLA10 showed that single mutations (I33E, L36E, or M43E) in the first helix were enough to impair dimerization (Maekawa et al., 2011). For RPM1 (RESISTANCE TO PSEUDOMONAS SYRINGAE PV. MACULICOLA 1), mutation of M34 or M41 weakened dimerization of the CC domain in coimmunoprecipitation assays, and mutation of 3 hydrophobic residues (I31E/M34E/M41E) was necessary to prevent CC dimerization in BiFC and coimmunoprecipitation assays (El Kasmi et al., 2017). It would be interesting to test the role of these residues in dimerization as ZAR1 contains hydrophobic residues at similar positions (Figure S1A).

All of the CC mutants as YFP fusions lost the ability to autoactivate immunity (Figure 2A, 2B). Since these mutations targeted different regions of the CC, these data indicate that the CC-YFP autoactivation phenotype is very sensitive to small disruptions and the overall folding of the domain. We therefore also tested the CC mutants in the context of full-length ZAR1, since this assay can identify weak effects on ZAR1 function. We found that ZAR1^H1^ retained some complementation of *NbZAR1*-VIGS, while the other mutants (ZAR1^H2a^, ZAR1^H2b^ and ZAR1^ED^) could not complement *NbZAR1*-VIGS. These data suggest that the α1a-α1b and α2-α3 loops (mutated in ZAR1^CCH1^ and ZAR1^H1^) only make a partial contribution to the overall folding of ZAR1^CC^. In addition, the α1a helix (mutated in ZAR1^CCH2a^, ZAR1^H2a^, ZAR1^CCH2b^ and ZAR1^H2b^) is important for ZAR1 function. Interestingly, the EDVID motif in ZAR1^CC^ (mutated in ZAR1^CCED^ and ZAR1^ED^) partially contributed to homodimerization and was essential for the interaction with ZAR1^NBARC1^, as well as the activation of downstream signaling (Figures 1, 2, S1). The EDVID motif was first identified in Rx, an NLR that recognizes the coat protein of *Potato Virus X* (Bendahmane et al., 1999; Rairdan et al., 2008). In Rx, the EDVID motif contributes to intramolecular interactions between the CC and NBARC domains (Rairdan et al., 2008). In MLA10, the EDVID motif is surface-exposed and positioned at the dimer interface (Maekawa et al., 2011), suggesting that it contributes to CC homodimerization. RPM1, an NLR involved in the recognition of *P. syringae* effector AvrRpm1, also contains an EDVID motif that may contribute to the self-association of CC domains (El Kasmi et al., 2017). In agreement with previous studies, the EDVID motif in ZAR1 is predicted to be surface-exposed (Figure S1), and is involved in CC dimerization (Figure 1C) and in the CC-NBARC interaction (Figure 3C). An outstanding question in the field is how the CC domain of various NLRs can activate immunity. TIR domains in NLRs have commonly been thought of as scaffolding proteins (Vajjhala et al., 2017). However, TIR domains in bacteria and Archaea were recently shown to act as NADase enzymes (Essuman et al., 2018). CC domains have no homology to proteins with known enzymatic activity. Thus it remains unclear whether they act directly or indirectly as a signal for immunity. Our data further support that the CC domain acts a signal for immunity because ZAR1^CC^-YFP could still trigger a HR in *NbZAR1*-silenced plants (Figure S3).

We demonstrated that the CC and NBARC domains of ZAR1 interact (Figure 3A), and that the NB domain is sufficient for the interaction and for suppression of ZAR1^CC^-YFP autoactivity (Figure 3D). Interactions between the CC and NBARC domain have also been observed for RPM1 (El Kasmi et al., 2017) which recognizes the *P. syringae* effector protein AvrRpm1, RPS5 (Ade et al., 2007) which recognizes the *P. syringae* effector protein AvrPphB, and the autoactive Rp1 NLR in maize (Wang et al., 2015a). Interestingly, the ZAR1^CCH2a^ and ZAR1^CCH2b^ constructs, that carried nine mutations in the first helix, were still able to interact very weakly with ZAR^NBARC1+2^. In contrast, only 2 mutations in the first helix (Y3A/M10A) of Rx were enough to completely disrupt the interaction between Rx^CC^ and Rx^NB-ARC1+2-LRR^, and the V6E/L9E/I13E mutations severely reduced the interaction (Slootweg et al., 2018). Unlike ZAR1, the CC-NBARC interaction in Rx appears to be sufficiently rigid that expression of either half of the CC domain of Rx1 (i.e. α1a-α1b or α2-α3) with Rx1^NB-ARC1+2-LRR^ led to the reconstitution of a functional Rx1 (Slootweg et al., 2018).

We sought to take advantage of loss-of-function mutations in ZAR1^NBARC1+2^ to identify whether they contributed to the CC-NBARC intramolecular interaction. However, these five mutations did not have any effect (Figure 4B). Consistent with this, the introduction of these mutations in a ZAR1^CCNBARC^-YFP variant did not restore the autoactivity observed in ZAR1^CC^-YFP. In Rp1 (Wang et al., 2015a), mutations that led to the loss of the CC-NBARC interaction in ZAR1 resulted in autoimmunity. Our loss-of-function mutations are found in or close to motifs that are believed to be particularly important for nucleotide binding and ATP hydrolysis (Collier and Moffett, 2009), including the P-loop (K195 and V202), the RNBS-B/kinase3 motif (S291) and the GLPL motif (P359) (Figure 6A, S7). Given the contribution of these residues to resistance, we speculate that they are critical for NBARC function in the activation of ZAR1 by effectors. Since our predicted autoactive mutations did not confer autoactivity (Figure S7), we were unable to test whether ZAR1 intramolecular or intermolecular interactions would be affected in the active state of the NLR.

We further demonstrated that the NBARC and LRR domains of ZAR1 interact in planta. The interaction was stronger with ZAR1^NB^ alone but could still be observed with the full-length NBARC domain (Figure 5A). Interactions between the NBARC and LRR domains have also been observed for RPS5 and RPP1 (Ade et al., 2007; Schreiber et al., 2016b), and for RPS5 this interaction maintains the inactive state (Qi et al., 2012). Interestingly, for Rx1, an acidic loop in the ARC2 domain is particularly important for interaction with a basic patch in the N-terminal region of the LRR domain (Slootweg et al., 2013). However, ZAR1 behaves more like RPP1 (Schreiber et al., 2016b) than Rx1, because the full NBARC domain containing the ARC2 subdomain exhibits a weaker interaction with the LRR domain. As seen with the CC-NBARC domain interactions (Figure 4B), mutations in the NBARC domain did not affect the interaction with LRR domain (Figure 5C).

Structural analysis of the LRR domain of ZAR1 identified 13 LRRs, followed by an extended region of the LRR domain, that could form another LRR segment or two helices (Figure 6B, S8). We found that loss-of-function mutations in the LRR domain had no effect on the interaction with the NBARC domain (Figure 5D). We previously showed that G645 and P816, but not S831, are necessary for the interaction between ZAR1^LRR^ and ZED1 (Baudin et al., 2017). The S831 mutation is located in the LRR extension (Figure 6B, S8). When we deleted this region (in ZAR1^Δ1^ containing aa1-824), ZAR1^Δ1^ was no longer able to complement *NbZAR1*-VIGS (Baudin et al., 2017). While *zar1*^S831F^ lacks an immune response to HopZ1a, ZAR1^LRR S831F^ can still interact with ZED1 (Baudin et al., 2017), and with ZAR1^NBARC1+2^ (Figure 5D). These data suggest that the LRR extension plays a distinct but unknown role in ZAR1 function and immunity.

Despite efforts by many groups to understand the dynamics and regulation of plant NLRs (Bonardi and Dangl, 2012), there is still a broad diversity of protein-protein interactions and higher-order macromolecular complexes that are critical for immunity. Oligomerization of NLRs is common for mammalian NLRs (Lechtenberg et al., 2014), and has only been rarely demonstrated for plant NLRs (Gutierrez et al., 2010). We propose the following model for ZAR1 interactions and activation (Figure 7). Inactive ZAR1 complexes are kept in an “off” state by intramolecular interactions, and intermolecular interactions with ZED1. We speculate that ZAR1 exists in an equilibrium between ZAR1 alone and ZAR1-ZED1 or ZAR1-ZRK complexes. This would provide sufficient plasticity for ZAR1 to exist as pre-activation complexes with different ZRKs, allowing ZAR1 to “switch” between different RLCKs, and provide specificity for different effector proteins. According to our model and experimental data, the inactive ZAR1 is tightly folded with both the LRR and CC domain interacting with the central NBARC domain (Figure 7). We previously showed that both the CC and LRR domain of ZAR1 interact with ZED1 in the absence of pathogens (Baudin et al., 2017). However, our data do not show whether both domains interact with ZED1 at the same time, and how ZED1 impacts ZAR1 intramolecular interactions. We propose that modification of the guardee by the effector induces a series of conformational rearrangements, which allows binding of ATP by the NBARC domain and dimerization of ZAR1^CC^. However, dimerization of ZAR1^CC^ is not sufficient for autoactivity, indicating that dimerization precedes autoactivity and the CC domain is more likely to act as a signal. Although the lack of autoactive mutations for ZAR1 does not allow us to specifically analyze the intramolecular interactions present in active complexes, we demonstrated that interactions between the domains must have a certain versatility to allow for conformational changes upon activation. Our data show that ZAR1 is regulated by subtle conformational changes that are finely controlled, and that ZAR1 displays specificity in its intramolecular interactions despite broad structural similarity with other NLRs. The ZAR1-ZED1 system demonstrates the complex and intricate ways in which immune receptors are dynamically regulated in vivo.

**Figure 7.**
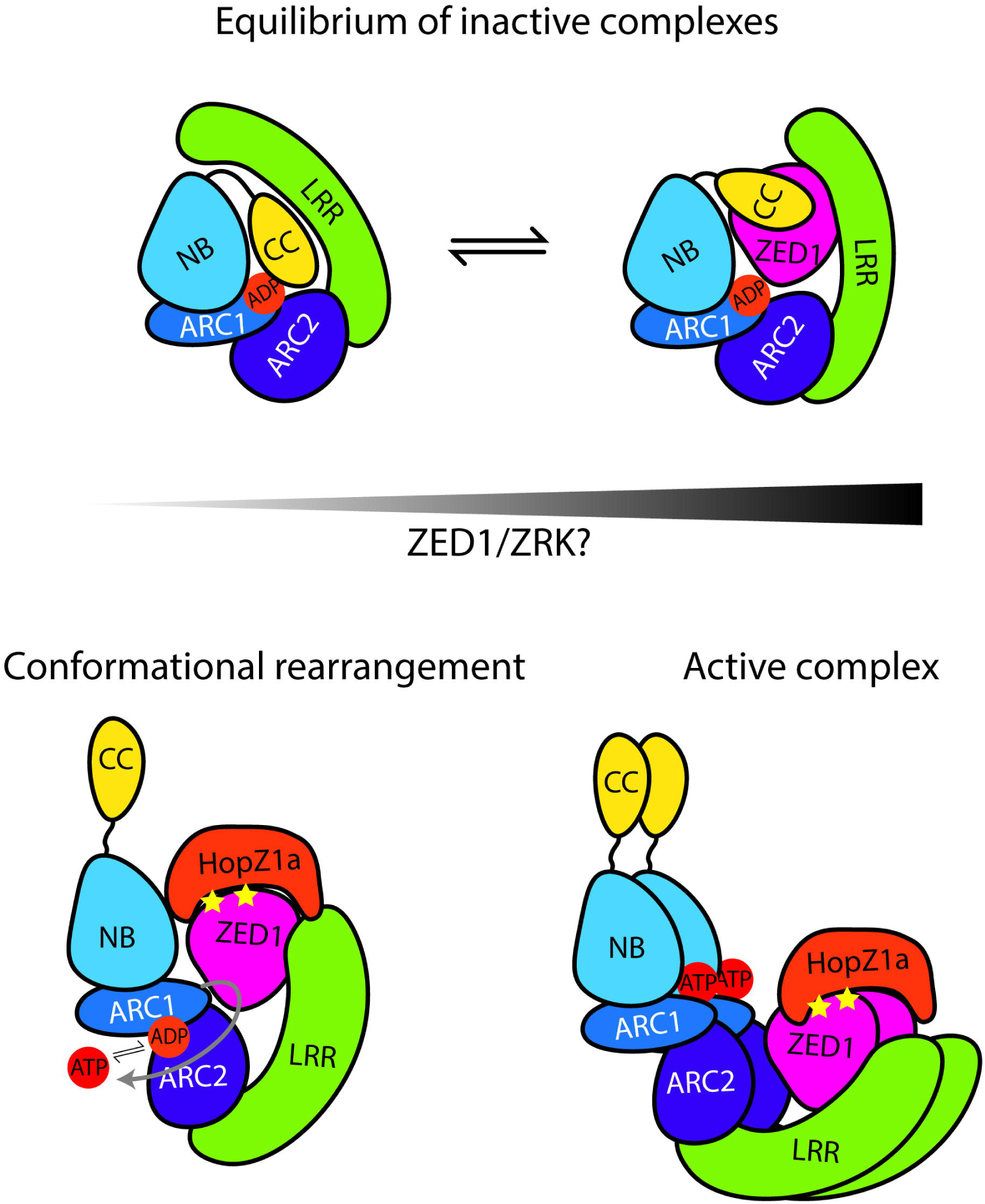
Model for ZAR1 activation. In the absence of the pathogen, ZAR1 is present in an equilibrium between inactive complexes lacking or containing ZED1 or ZRKs. We propose that inactive ZAR1 versus inactive ZAR1/ZED1/ZRKs complexes would be necessary to allow for interactions of ZAR1 with diverse RLCKs. We propose that acetylation of ZED1 (shown with stars) by HopZ1a leads to conformational changes in ZAR1 that make the CC domain more accessible, allow the exchange of ADP for ATP, and cause rotation of the ARC2 domain. Although dimerization of ZAR1^CC^ has been shown, it is unknown whether the rest of the protein also self-associates. Intramolecular and intermolecular conformational changes trigger activation of ZAR1 and ETI.

## MATERIALS AND METHODS

### Computational analysis and 3D modeling

The CC, NBARC and LRR domains of ZAR1 were modelled individually by remote homology modeling workflows similar to those used to model equivalent domains in Lr10 (LEAF RUST 10 from *Triticum dicoccoides* [emmer wheat]), NRC1 (NB-LRR REQUIRED FOR HYPERSENSITIVE RESPONSE-ASSOCIATED CELL DEATH 1 from *Solanum lycopersicum* [tomato]), Rx and Pm3 (POWDERY MILDEW 3 from *Triticum aestivum*) (Sela et al., 2012; Slootweg et al., 2013; Sela et al., 2014; Sueldo et al., 2015). In brief, profiles were raised for sequence variability and sequence propensities for secondary structure, intrinsic disorder, accessibility, linkers, coil-coil and turn local structure. For each profile, several methods were used and the consensus was built to increase the prediction reliability as previously described (Sela et al., 2012; Slootweg et al., 2013; Sela et al., 2014; Sueldo et al., 2015). These structural propensities were further used to refine sequence alignments with the closest domain templates in each case. Alignments were generated by using appropriate BLOSUM matrices with either the EMBOSS-stretcher sequence-only pair-wise alignment method (Myers and Miller, 1988) for single template models or PRALINE multiple sequence alignment method enhanced by incorporating PSSM profiles and secondary structure predictions (Bawono and Heringa, 2014) when dealing with multiple potential structural templates.

ZAR1^CC^ was modeled in both the monomeric 4-bundle helix configuration (1CC4α) and the 2-helix dimer conformation (2CC2α). The 1CC4α model was built starting from the Sr33 NMR structure (PDB: 2NCG, (Casey et al., 2016)) and the Rx crystal structure (PDB: 4M70, (Hao et al., 2013)) with which ZAR1 shares 17% and 15% sequence identity respectively. The 2CC2α model was built from the dimer configuration of MLA10 structures (PBD: 3QFL, (Maekawa et al., 2011)) with which ZAR1-CC sequence shares 19% identity.

ZAR1^NBARC^ was modeled in both closed and opened conformations starting from human Apaf1 structures in the closed ADP-bound state (PDB: 1Z6T, (Riedl et al., 2005)) and open ATP-bound state (PDB: 5JUY, (Cheng et al., 2016)) respectively. We used Apaf1, as ZAR1 shares 20% sequence identity with Apaf1, which was the best score among all experimental NBARC structures.

ZAR1^LRR^ was modeled using the Optimized Joined Fragment Remote Homology modeling (OJFRHM) method previously used to model the LRR domains of Lr10, Rx and Pm3 (Sela et al., 2012; Slootweg et al., 2013; Sela et al., 2014). From our structural LRR database, the closest local ZAR1 templates were PBD: 2RA8, 1OGQ and 4U06 (Vorobiev et al., 2007; Di Matteo et al., 2003; Miras et al., 2015).

All homology models were built with Discovery Studio software from Accelrys-Dassault Systemes and Modeller v9.17 (Webb and Sali, 2014). Within the sequence conserved regions (SCR), the model was raised by coordinate transfer and side chain rebuilding, while the sequence variable regions (SVRs) were randomly generated and filtered by steric constraints, followed by successive rounds of energy minimization (EM) and simulated annealing (SA). The global model generated as above was further subjected to overall structural optimization by repeated rounds of EM, SA and Molecular Dynamics in explicit solvent performed with CHARMM36 force field in NAMD (Phillips et al., 2005), followed iteratively by model quality check until convergence. Model quality was assessed with MolProbity (Davis et al., 2007) and PROCHECK V.3.5 (Laskowski et al., 1993) for crystallographic standards compliance. Trajectory analysis was performed using VMD (Humphrey et al., 1996) and all figures of predictive models’ structures were created using PyMOL Molecular Graphics System v1.3.

### Plasmid construction

Phusion polymerase (New England Biolabs) was used for all cloning, and all constructs were confirmed by sequencing. Sequence analysis was performed with CLC Main Workbench. For the VIGS constructs, we used a clone carrying a genomic fragment of *NbZAR1*, as described in Baudin et al., 2017. The single mutations were generated using the Q5 Site-directed mutagenesis kit (New England Biolabs). For ZAR1^CCH1^, ZAR1^CCH2a^ ZAR1^CCH2b^ and ZAR1^CCED^, the gene fragments were synthesized as gBlocks by Integrated DNA Technologies, and then used as templates for PCR. For the complementation assay, the ZAR1^CC^ mutations were introduced in the full-length gene using the internal restriction site *EcoRI*. ZAR1 variants were amplified by PCR to add a 5’*XhoI* site and cloned into the pMac15 vector to maintain the frame for the vector-encoded C-terminal 5xMyc tag. The pMac15 vector was modified from pBD to contain a 5xMyc tag between the *StuI* and *SpeI* sites, as described in Baudin et al., 2017. The ZED1 3xFlag construct was cloned by a crossover PCR approach to add the 3xFlag tag in frame at the 3’end. The PCR products were then cloned into the pMac14 vector, which was modified from pBD (Aoyama and Chua, 1997; Lewis et al., 2008). The 3xFlag tagged NBARC variants were amplified by PCR to add a 5’*XhoI* site and cloned into the pMac16 vector to maintain the frame for the vector-encoded C-terminal 3xFlag tag. The pMac16 vector was modified from pBD to contain a 3xFlag tag between the *StuI* and *SpeI* sites. For the split YFP and YFP fusions, the genes were amplified by PCR to contain a 5’*XhoI* restriction site. The 3’ primers were designed to maintain the reading frame of the C-terminal fusion. pBD-YFP, pBD-nYFP, and pBD-cYFP were modified from pTA7002 (Aoyama and Chua, 1997) to add an HA tag and full-length YFP, nYFP (residues 1-155), or cYFP (residues 156 to the stop codon) between the *StuI* and *SpeI* sites (Lewis et al., 2014).

For protein interaction analyses in yeast, all constructs were prepared in the DupLEX-A system (OriGene Technologies, Inc., Rockford, MD) using the vectors pEG202 and pJG4-5 to express bait and prey proteins, respectively. All sequences were cloned into the *Eco*RI and *Xho*I sites of these vectors. to maintain the frame of the vector-encoded fusions. The pJG4-5 vector contains a nuclear localization signal, B42 activation domain and HA tag at the 5’ end. The pEG202 vector contains the LexA DNA binding domain at the 5’ end.

### Yeast two-hybrid experiments

All constructs were transformed into yeast using Frozen-EZ Yeast Transformation II reagents (Zymo Research Corp., Irvine, CA) to introduce pEG202 constructs into yeast strain RFY206 (mating type A), while pJG4-5 constructs were introduced into strain EGY48 (mating type α). Transformants were recovered on Synthetic Defined (SD)+Glucose media lacking histidine and uracil to select for colonies containing pEG202 constructs, or lacking tryptophan to select for pJG4-5 transformants. Yeast matings were performed in YPDA media at 30°C overnight, followed by two rounds of diploid selection on SD+Glucose-HisUraTrp plates at 30°C overnight. To assay for protein-protein interactions, diploid yeast were first resuspended in 150 μL of sterile distilled water and adjusted to an OD_600_ of 2, followed by three serial 1:5 dilutions in water. Five microliters of each dilution were spotted on SD+Glucose-HisUraTrpLeu and SD+Galactose+Raffinose-HisUraTrpLeu plates, with yeast growth indicating the activation of a LEU2 reporter. Images of yeast plates were generally captured after three days of growth at 30°C.

### Coimmunoprecipitation

In planta coimmunopurification was performed using 5 cm^2^ of *N. benthamiana* leaves transiently expressing our genes of interest. Tissue was homogenized in liquid nitrogen, and protein complexes were extracted using 1 mL of IP1 buffer (50 mM HEPES, 50 mM NaCl, 10 mM EDTA, 0.2% TritonX-100, and 0.1 mg/mL Dextran [Sigma-Aldrich D1037], pH 7.5). The clarified extract was then mixed with 10 µL of anti-Flag magnetic beads [Sigma-Aldrich M8823] for 3 h at 4°C. The coimmunoprecipitated proteins were washed four times with IP2 buffer (50 mM HEPES, 150 mM NaCl, 10 mM EDTA, and 0.1% Triton X-100, pH 7.5) on a magnetic stand. IP1 and IP2 buffers were supplemented with proteinase inhibitor cocktail (1 mM PMSF, 1 mg/mL leupeptin [Sigma-Aldrich L2023], 1 mg/mL aprotinin [Sigma-Aldrich A6191], 1 mg/mL antipain [Sigma-Aldrich A1153], 1 mg/mL chymostatin [Sigma-Aldrich C7268], and 1 mg/mL pepstatin [Sigma-Aldrich P5315]). The immunopurified fraction was eluted incubating the beads with 30 µL of 150 µg/mL 3xFlag peptide [Sigma-Aldrich F4799] for 30 minutes at room temperature. Finally, the input and immunopurified fractions were separated on 10% acrylamide gels by SDS-PAGE and detected by immunoblot using α-c-Myc antibody (Abcam ab32072), or α-Flag antibody (Sigma-Aldrich F1804) followed by the appropriate secondary antibody coupled with horseradish peroxidase.

### Plant material and transient expression

We followed protocols previously described in Baudin et al., 2017 with some slight modifications as follows. *Nicotiana benthamiana* plants were grown in a growth chamber at 22°C with 9 h light (∼130 µEm^−2^s^−1^) and 15 h dark cycles. The seeds were sterilized in 10% bleach for 10 min and then washed ten times with sterile nanopure water. After germination on soil, seedlings were transplanted onto individual pots and used 3 to 5 weeks after transplanting. For the *Agrobacterium*-mediated transient expression, *Agrobacterium tumefaciens* GV2260 cultures were grown overnight at 28°C in LB broth with kanamycin and rifampicin. The next day, the bacteria were resuspended in 1 mL of 10 mM MES, pH 5.6 and 200 µM of acetosyringone and incubated in the dark for ∼4 h. The cultures were then adjusted to an optical density (λ=600 nm) of 1.0. The cultures containing each plasmid were mixed in equal volumes to a final optical density of 0.25 per construct. For BiFC experiments, *A. tumefaciens* carrying the gene silencing suppressor P19 was co-infiltrated with the constructs at an optical density of 0.25 (Voinnet et al., 2003). The underside of the leaves of 5- to 7-week-old *N. benthamiana* plants were infiltrated by hand with a needleless syringe. For dexamethasone-inducible constructs, the plants were sprayed with 20 µM dexamethasone (Sigma-Aldrich) 12 to 24 h after inoculation. Tissues were collected from 3 to 48 h after dexamethasone induction depending on the experiment. For the VIGS experiments, the plants were used 1 to 2 weeks after inoculation.

### Phenotyping and conductivity

To phenotype the HR-like phenotype induced by ZAR1^CC^-YFP autoactivity, infiltrated leaves were observed ∼40 h after dexamethasone induction. The severity of HR was rated on the following phenotypic scale: no HR, weak HR, medium HR and strong HR. For ion leakage assays in *N. benthamiana*, two disks (1.5 cm^2^) were harvested 20 h after infiltration (3 h after dexamethasone induction) floated in nanopure water for 1 h and transferred to 6 mL of nanopure water. Readings were taken with an Orion 3 Star conductivity meter (Thermo Electron) at 24 h (T0) and 48 h (T24) after infiltration.

### Confocal microscopy

*N. benthamiana* leaves transiently expressing the constructs of interest were infiltrated with water and mounted on microscope slides. Samples were imaged using a Leica SP8 confocal laser-scanning microscope equipped with a 40x water-immersion objective (HC PL APO CS2 40x/1.10 WATER). The 514 nm argon laser line was used to excite YFP, and florescence was observed using the specific emission window of 520 to 600 nm. The laser power, gain, zoom and average settings were kept consistent over the same image series to allow fluorescence intensity comparison across samples. Images were processed using the Leica Application Suite X software.

### Western blots

1 cm2 of leaf tissue was collected 24 h after dexamethasone induction and frozen in liquid nitrogen. The frozen tissue was then homogenized and proteins were extracted in 100 µL of 1X Laemmli buffer and boiled for 5 min. Proteins were then separated on 10% acrylamide SDS-PAGE gels. After transferring onto nitrocellulose membranes, tagged proteins were detected using α-c-Myc antibody (Abcam ab32072), α-GFP antibody (Roche 11814460001) or α-Flag antibody (Sigma-Aldrich F1804), followed by the appropriate secondary antibody coupled to horseradish peroxidase. Protein expression in yeast was evaluated following shaking incubation of cultures at 30°C overnight in SD+Glucose-HisUra (pEG202 constructs) or SD+Galactose+Raffinose-Trp (pJG4-5) media. Cultures were processed for protein extraction according to a lithium acetate/sodium hydroxide pre-treatment protocol described by (Zhang et al., 2011). Samples were resolved on 8% discontinuous SDS-PAGE gels, transferred to nitrocellulose membranes, and detected by either α-HA (Roche 12CA5) or α-LexA antibodies (Santa Cruz Biotechnology SC7544).

### Statistical analyses

The conductivity data were analyzed using Minitab 17 software (Minitab). For each conductivity assay, the T24 data were analyzed by a one-way ANOVA statistical test at a significance level of p = 0.05 followed by a multiple comparisons of means using Tukey’s post hoc test.

### Accession numbers

Sequence data from this article can be found in the GenBank/EMBL data libraries under accession numbers At3g50950 (AtZAR1), At3g57750 (AtZED1), and Niben101Scf17398g00012 (NbZAR1).

## Supplemental Data

The following supplemental materials are available.

Supplemental Figure S1. Alignment and modeling of ZAR1^CC^ and CC domain mutants.

Supplemental Figure S2. Protein expression in yeast strains used for two-hybrid analyses.

Supplemental Figure S3. ZAR^CC^ autoactivity in *N. benthamiana* is independent from *NbZAR1*.

Supplemental Figure S4. BiFC analysis of the ZAR1^CC^ – ZAR1^NBARC^ interaction and protein expression of ZAR1 domain constructs.

Supplemental Figure S5. Residues associated with autoactive NLRs do not activate immunity in ZAR1.

Supplemental Figure S6. The NBARC and LRR domains interact in planta by bimolecular fluorescence complementation.

Supplemental Figure S7. ZAR1^NBARC^ alignments used to build the closed and open conformation of ZAR1^NBARC^ 3D models starting from human Apaf1 crystal structures.

Supplemental Figure S8. ZAR1^LRR^ alignments and predicted/crystal secondary structure used in modeling the LRR domain.

## ACKNOWLEDGMENTS

We thank Jana A. Hassan from the Lewis Lab for providing constructive comments and suggestions on the manuscript.

## Author contributions

MB, ECM, AJP, KJS and JDL designed research. MB performed *in planta* experiments. KJS performed yeast experiments. ECM and AJP contributed new analytic/computational tools. MB, ECM, AJP, KJS and JDL analyzed data. MB, AJP, KJS and JDL wrote the paper.

## SUPPLEMENTAL FIGURE LEGENDS

**Supplemental Figure S1.** Alignment and modeling of ZAR1^CC^.

A, The Sr33 and Rx structures were used to generate the 1CC4α monomeric model, while MLA10 structure was used to generate the 2CC2α dimer model of ZAR1^CC^. The alignment of ZAR1^CC^ to RPM1 is also shown. Conserved hydrophobic residues are shown in black boxes, and the EDVID motif is shown in a brown box. Amino acids are colored based on their properties as follows: hydrophobic (yellow), positively charged (blue), negatively charged (red), proline and glycine (green), and polar (grey).

B, Schematic (representation of a four-helices bundle monomer, a two-helices bundle monomer and a hairpin dimer. The different helical segments are indicated as α1a, α1b, α2 and α3. The 4 consecutive helices are colored from bright orange to dark red. N indicates the N-terminus of the CC domain and C indicates the C-terminus of the CC domain.

C, Mutations were mapped onto the 3D models 1CC4α (CC monomer, left) and 2CC2α (CC dimer, right) respectively. The 4 consecutive helices are colored as in Figure 1A and 1B, from bright orange to dark red. The region corresponding to the first turn in monomeric 1CC4α, which breaks the long α1 helix of the dimer in α1a & α1b, is shown in green. Mutated amino acids are shown with side chains in sticks&dots representation.

**Supplemental Figure S2.** Protein expression in yeast strains used for two-hybrid analyses. Yeast cultures were incubated with shaking at 30°C overnight in SD+Glucose-HisUra (for pEG202 constructs) or SD+Galactose+Raffinose-Trp (for pJG4-5 constructs media. Proteins expressed from pEG202 were detected with an α -LexA antibody, while an α -HA antibody was used to detect proteins expressed from pJG4-5.

**Supplemental Figure S3.** ZAR^CC^ autoactivity in *N. benthamiana* is independent from *NbZAR1*. *A. tumefaciens* carrying constructs expressing ZAR1^CC^-YFP, HopZ1a + ZED1, HopZ1a + ZED1 + ZAR1 or Control-YFP (At3g46600) were infiltrated into *N. benthamiana* leaves silenced for *GUS* or *NbZAR1* genes. The HR is shown 48 h after dexamethasone induction (top). The scale bar is 1 cm. The graph (bottom) quantitates the HR strength for each construct. The experiment was repeated three times with similar results.

**Supplemental Figure S4.** BiFC analysis of the ZAR1^CC^ – ZAR1^NBARC^ interaction and protein expression of constructs.

A, Bimolecular fluorescence complementation of ZAR1^CC^ and ZAR1^NBARC^ interactions. Equal amounts of *A. tumefaciens* carrying ZAR1^CC^ or empty vector (EV) as a fusion to cYFP, and *A. tumefaciens* carrying ZAR1^NBARC^ truncations as a fusion to nYFP, were mixed and infiltrated. Leaf sections were imaged 24 h after dexamethasone induction using a Leica SP8 confocal scanning microscope. The YFP channel is shown for all images. The scale bar is 50 µm. The experiment was repeated 3 times with similar results.

B, Immunoblot analysis of ZAR1, ZAR1 truncations or control (At3g46600) proteins as fusions to YFP. Samples were harvested 24 h after dexamethasone induction. Equal amounts of proteins were resolved on 10% SDS-PAGE gels, blotted onto nitrocellulose, and probed with GFP antibodies in Western blot analysis (WB). The Ponceau red stained blot was used as the loading control.

C, Immunoblot analysis of ZAR1^CC^, ZAR1 truncations carrying mutations, or control (At3g46600) proteins as fusions to YFP, as in B.

**Supplemental Figure S5.** Residues associated with autoactive NLRs do not activate immunity in ZAR1.

A, Schematic representation of putative autoactive mutations in ZAR1. Residues in located in the following motifs as follows: G194A/K195A in the P-loop, D268E in the kinase 2 motif, and D489V in the MHD motif. B, HR assay of putative ZAR1 autoactivation mutants. *A. tumefaciens* carrying wild type ZAR1 with a myc tag, ZAR1 mutants with a myc tag, or empty vector (EV) were pressure-infiltrated into *N. benthamiana* leaves. *A. tumefaciens* carrying HopZ1a with a HA tag, and *A. tumefaciens* carrying ZED1 with a flag tag, were mixed to the same optical density and pressure-infiltrated into *N. benthamiana* leaves. Constructs were expressed under a dexamethasone-inducible promoter. Photographs were taken 48 h after dexamethasone application. The experiment was repeated 3 times with similar results. C, Immunoblot analysis of ZAR1 or ZAR1 mutant proteins as fusions to the Myc tag. Samples were harvested 24 h after dexamethasone induction. Equal amounts of proteins were resolved on 10% SDS-PAGE gels, blotted onto nitrocellulose, and probed with Myc antibodies in Western blot analysis (WB). The Ponceau red stained blot was used as the loading control.

**Supplemental Figure S6.** The NBARC and LRR domains interact in planta by bimolecular fluorescence complementation.

Equal amounts of *A. tumefaciens* carrying ZAR1^NBARC^ truncations or mutants, or empty vector (EV) as a fusion to cYFP, and *A. tumefaciens* carrying ZAR1^LRR^ as a fusion to nYFP, were mixed and pressure-infiltrated into *N. benthamiana* leaves. Constructs were expressed under a dexamethasone-inducible promoter. Leaf sections were imaged 24 h after dexamethasone induction using a Leica SP8 confocal scanning microscope. The YFP channel is shown for all images. The scale bar is 50 µm. The colored boxes correspond to the following domains: medium blue is the nucleotide-binding (NB) subdomain, dark blue is the Apaf1-R protein-CED4 1 (ARC1) subdomain, purple is the ARC2 subdomain, and green is the leucine-rich repeat (LRR) domain. NBARC1 indicates that the NB and ARC1 subdomains are present. NBARC1+2 indicates that the NB, ARC1 and ARC2 subdomains are present. The experiment was repeated 3 times with similar results.

**Supplemental Figure S7.** ZAR1^NBARC^ alignments used to build the closed and open conformation of ZAR1^NBARC^ 3D models starting from human Apaf1 crystal structures. Despite the rearrangement of the ARC2 domain which is rotated with respect to NB and ARC1, the internal structure of each subdomain remains almost unchanged in the two conformational states. The locations of K195N, V202M, S291N, P359L and L465F are shown with magenta arrows. Amino acids are colored based on their properties as follows: hydrophobic (yellow), positively charged (blue), negatively charged (red), proline and glycine (green), and polar (grey).

**Supplemental Figure S8.** ZAR1^LRR^ alignments and predicted/crystal secondary structure used in modeling the LRR domain.

LRR motifs are shown in blue boxes. The locations of G645E, P816Q and S831F are shown with magenta arrows. Amino acids are colored based on their properties as follows: hydrophobic (yellow), positively charged (blue), negatively charged (red), proline and glycine (green), and polar (grey).

