## Supplemental Figures 1-8 for "Structure-function analysis of ZAR1 immune receptor reveals key molecular interactions for activity"

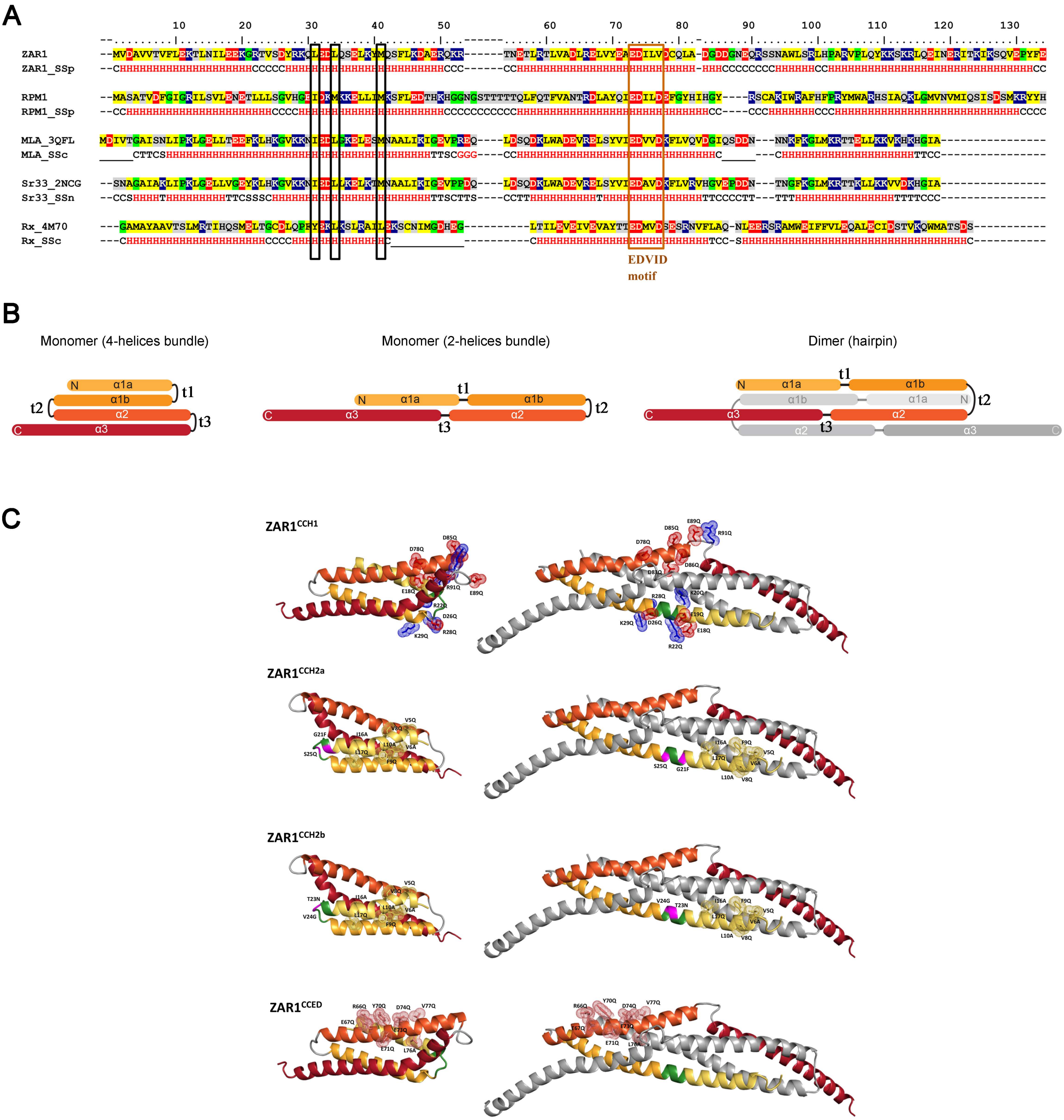


**Supplemental Figure S1. Alignment and modeling of ZAR1$^{CC}$**

A, The Sr33 and Rx structures were used to generate the 1CC4α monomeric model, while MLA10 structure was used to generate the 2CC2α dimer model of ZAR1$^{CC}$. The alignment of ZAR1$^{CC}$ to RPM1 is also shown. Conserved hydrophobic residues are shown in black boxes, and the EDVID motif is shown in a brown box. Amino acids are colored based on their properties as follows: hydrophobic (yellow), positively charged (blue), negatively charged (red), proline and glycine (green), and polar (grey). B, Schematic representation of a four-heli ces bundle monomer, a two-helices bundle monomer and a hairpin dimer. The different helical segments are indicated as α1a, α 1b, α2 and α3. The 4 consecutive helices are colored from bright orange to dark red, and the second molecule in the dimer is shown in grey. N indicates the N-terminus of the CC domain and C indicates the C-terminus of the CC domain. C, Mutations were mapped onto the 3D models 1CC4α (CC monomer - left) and 2CC2α (CC dimer - right) respectively. The 4 consecutive helices are colored as in Figure 1A and 1B, from bright orange to dark red. The region corresponding to the first turn in monomeric 1CC4α, which breaks the long α1 helix of the dimer in α1a & α1b, is shown in green. Mutated amino acids are shown with side chains in sticks&dots representation.

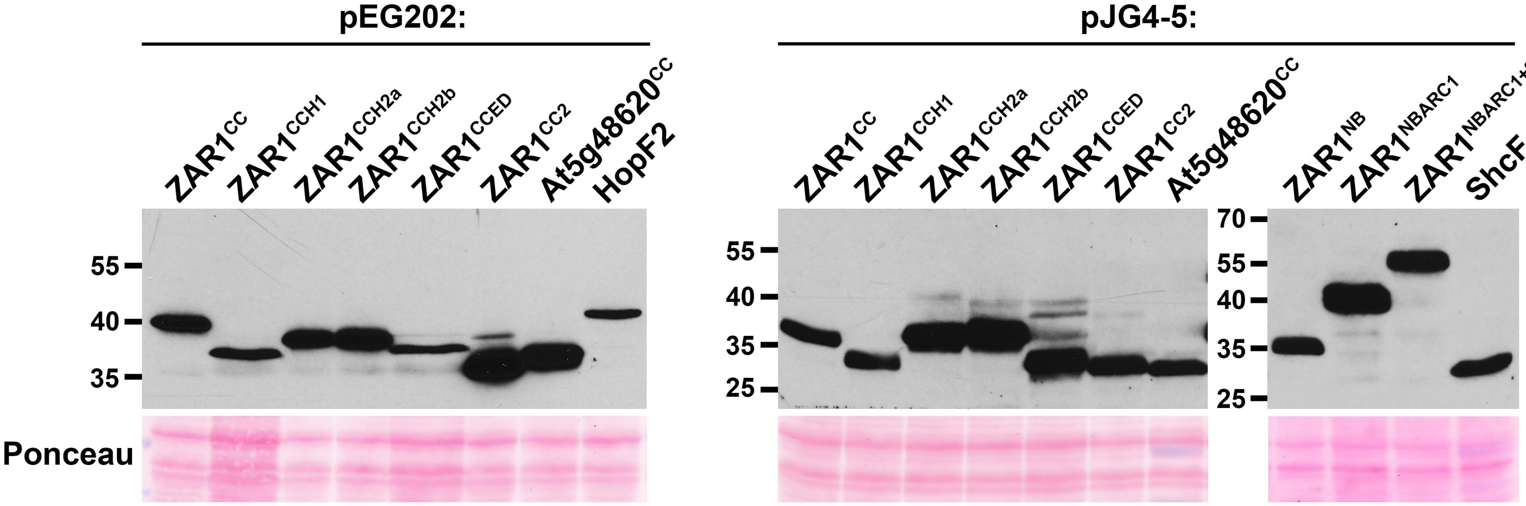


**Supplemental Figure S2.** Protein expression in yeast strains used for two-hybrid analyses. Yeast cultures were incubated with shaking at 30°C overnight in SD+Glucose-HisUra (pEG202 constructs) or SD+Galactose+Raffinose-Trp (pJG4-5) media. Proteins expressed from pEG202 were detected with an α -LexA antibody, while an α -HA antibody was used to detect proteins expressed from pJG4-5.

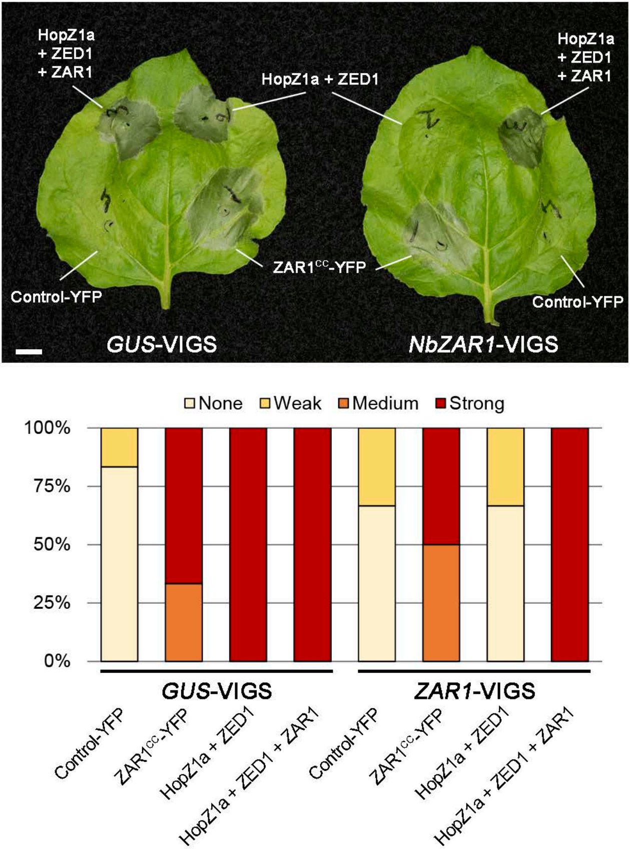


**Supplemental Figure S3. ZAR$^{CC}$ autoactivity in *N. benthamiana* is independent from *NbZAR1***

A *tumefaciens* carrying constructs expressing ZAR1$^{CC}$-YFP, HopZ1a + ZED1, HopZ1a + ZED1 + ZAR1 or Control-YFP (At3g46600) were infiltrated into *N. benthamiana* leaves silenced for GUS or NbZAR1 genes. The HR is shown 48 h after dexamethasone induction (top). The scale bar is 1 cm. The graph (bottom) quantitates the HR strength for each construct. The experiment was repeated three times with similar results.

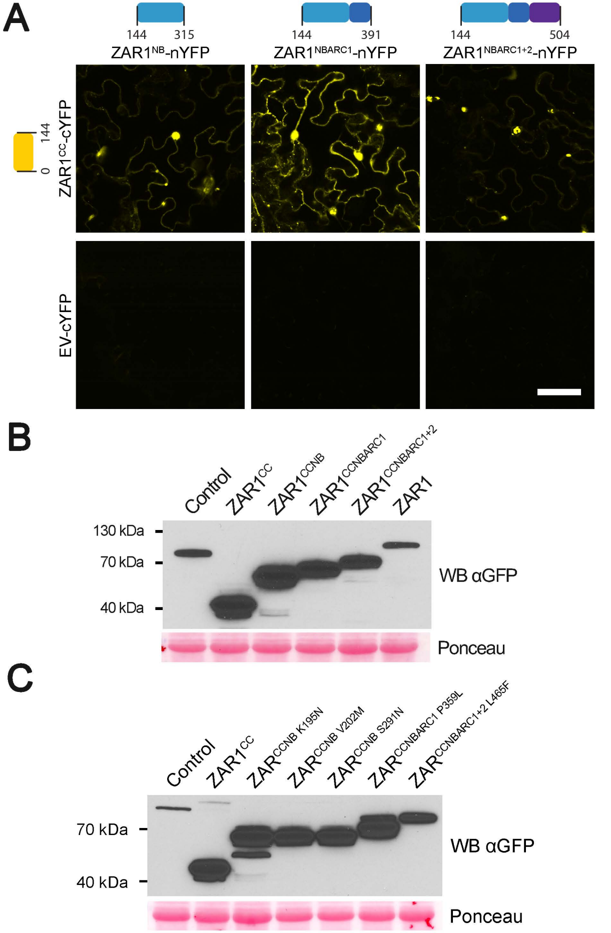


**Supplemental Figure S4. BiFC analysis of the ZAR1$^{CC}$ – ZAR1$^{NBARC}$ interaction and protein expression of constructs.** A, Bimolecular fluorescence complementation of ZAR1$^{CC}$ and ZAR1$^{NBARC}$ interactions. Equal amounts of *A. tumefaciens* carrying ZAR1$^{CC}$ or empty vector (EV) as a fusion to cYFP, and *A. tumefaciens* carrying ZAR1$^{NBARC}$ truncations as a fusion to nYFP, were mixed and infiltrated. Leaf sections were imaged 24 h after dexamethasone induction using a Leica SP8 confocal scanning microscope. The YFP channel is shown for all images. The scale bar is 50 µm. The experiment was repeated 3 times with similar results. B, Immunoblot analysis of ZAR1, ZAR1 truncations or control (At3g46600) proteins as fusions to YFP. Samples were harvested 24 h after dexamethasone induction. Equal amounts of proteins were resolved on 10% SDS-PAGE gels, blotted onto nitrocellulose, and probed with GFP antibodies in Western blot analysis (WB). The Ponceau red stained blot was used as the loading control. C, Immunoblot analysis of ZAR1$^{CC}$, ZAR1 truncations carrying mutations, or control (At3g46600) proteins as fusions to YFP, as in B.

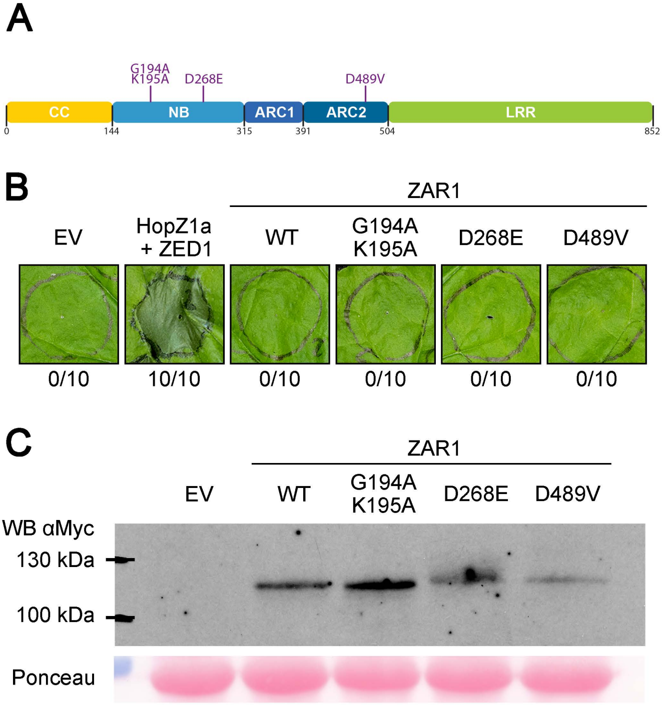


**Figure S5: Residues associated with autoactive NLRs do not activate immunity in ZAR1.**

(A) Schematic representation of putative autoactive mutations in ZAR1. Residues in located in the following motifs as follows: G194A/K195A in the P-loop, D268E in the kinase 2 domain, and D489V in the MHD motif.

(B) HR assay of putative ZAR1 autoactivation mutants. *A. tumefaciens* carrying wild type ZAR1 with a myc tag, ZAR1 mutants with a myc tag, or empty vector (EV) were pressure-infiltrated into *N. benthamiana* leaves. *A. tumefaciens* carrying HopZ1a with a HA tag, and *A. tumefaciens* carrying ZED1 with a flag tag, were mixed to the same optical density and pressure-infiltrated into *N. benthamiana* leaves. Constructs were expressed under a dexamethasone-inducible promoter. Photographs were taken 48 h after dexamethasone application. The experiment was repeated 3 times with similar results.

(C) Immunoblot analysis of ZAR1 or ZAR1 mutant proteins as fusions to the Myc tag. Samples were harvested 24 h after dexamethasone induction. Equal amounts of proteins were resolved on 10% SDS-PAGE gels, blotted onto nitrocellulose, and probed with Myc antibodies in Western blot analysis (WB). The Ponceau red stained blot was used as the loading control.

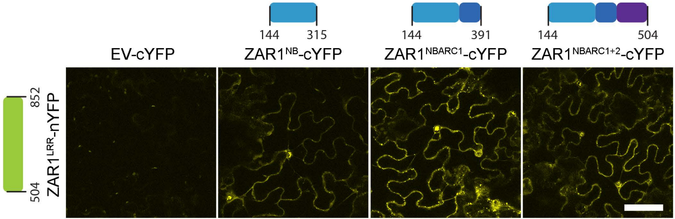


**Supplemental Figure S6.** The NBARC and LRR domains interact in planta by bimolecular fluorescence complementation. Equal amounts of *A. tumefaciens* carrying ZAR1$^{NBARC}$ truncations or mutants, or empty vector (EV) as a fusion to cYFP, and *A. tumefaciens* carrying ZAR1$^{LRR}$ as a fusion to nYFP, were mixed and pressure-infiltrated into *N. benthamiana* leaves. Constructs were expressed under a dexamethasone-inducible promoter. Leaf sections were imaged 24 h after dexamethasone induction using a Leica SP8 confocal scanning microscope. The YFP channel is shown for all images. The scale bar is 50 µm. The colored boxes correspond to the following domains: medium blue is the nucleotide-binding (NB) subdomain, dark blue is the Apaf1-R protein-CED4 1 (ARC1) subdomain, purple is the ARC2 subdomain, and green is the leucine-rich repeat (LRR) domain. NBARC1 indicates that the NB and ARC1 subdomains are present. NBARC1+2 indicates that the NB, ARC1 and ARC2 subdomains are present. The experiment was repeated 3 times with similar results.

### ZAR1_NBS

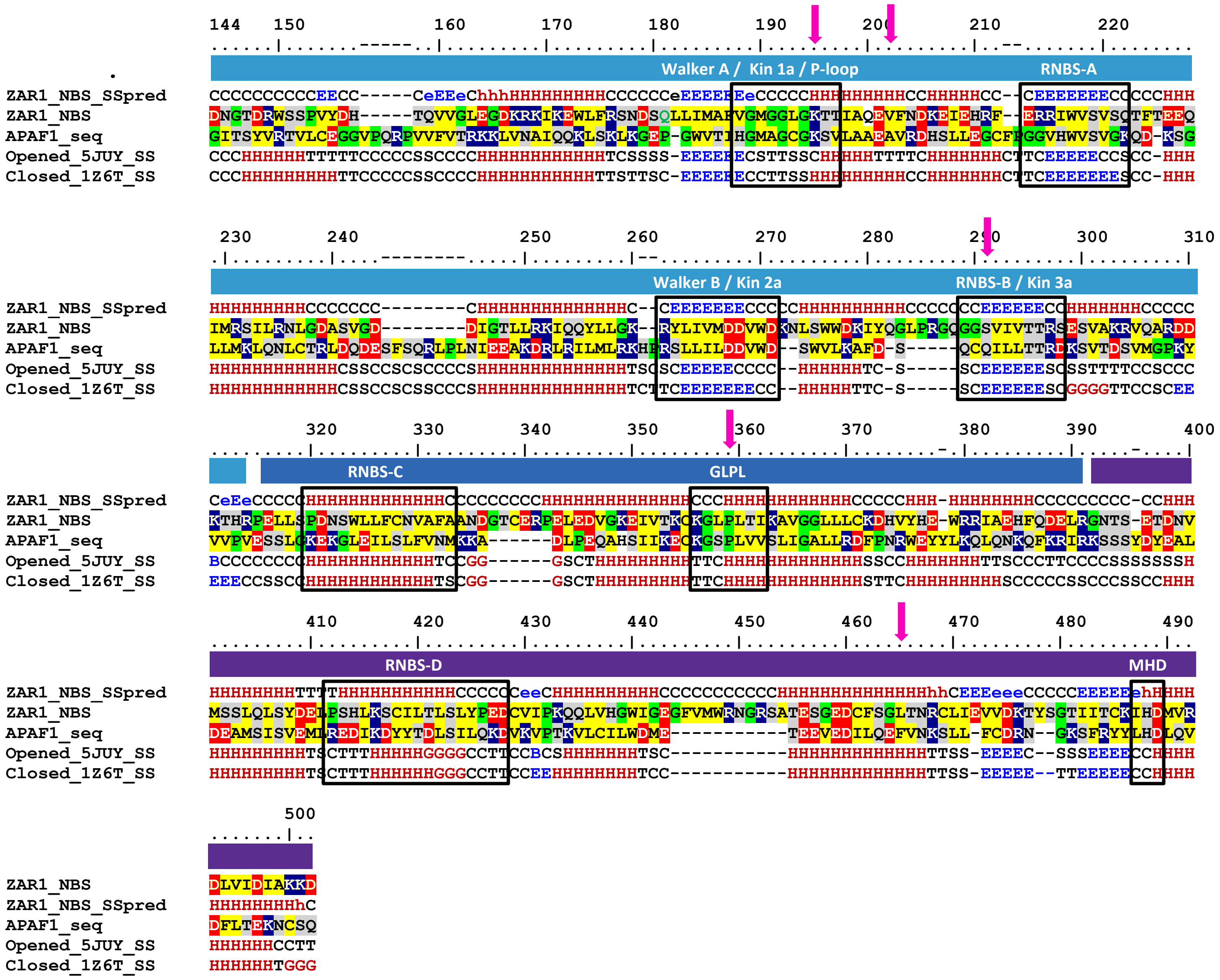


**Figure S7: ZAR1$^{NBARC}$ alignments used to build the closed and open conformation of ZAR1$^{NBARC}$ 3D models starting from human Apaf1 crystal structures**. Despite the rearrangement of the ARC2 domain which is rotated with respect to NB and ARC1, the internal structure of each subdomain remains almost unchanged in the two conformational states. The location of K195N, V202M, S291N, P359L and L465F are shown with magenta arrows. Amino acids are colored based on their properties as follows: hydrophobic (yellow), positively charged (blue), negatively charged (red), proline and glycine (green), and polar (grey).

ZAR1_LRR

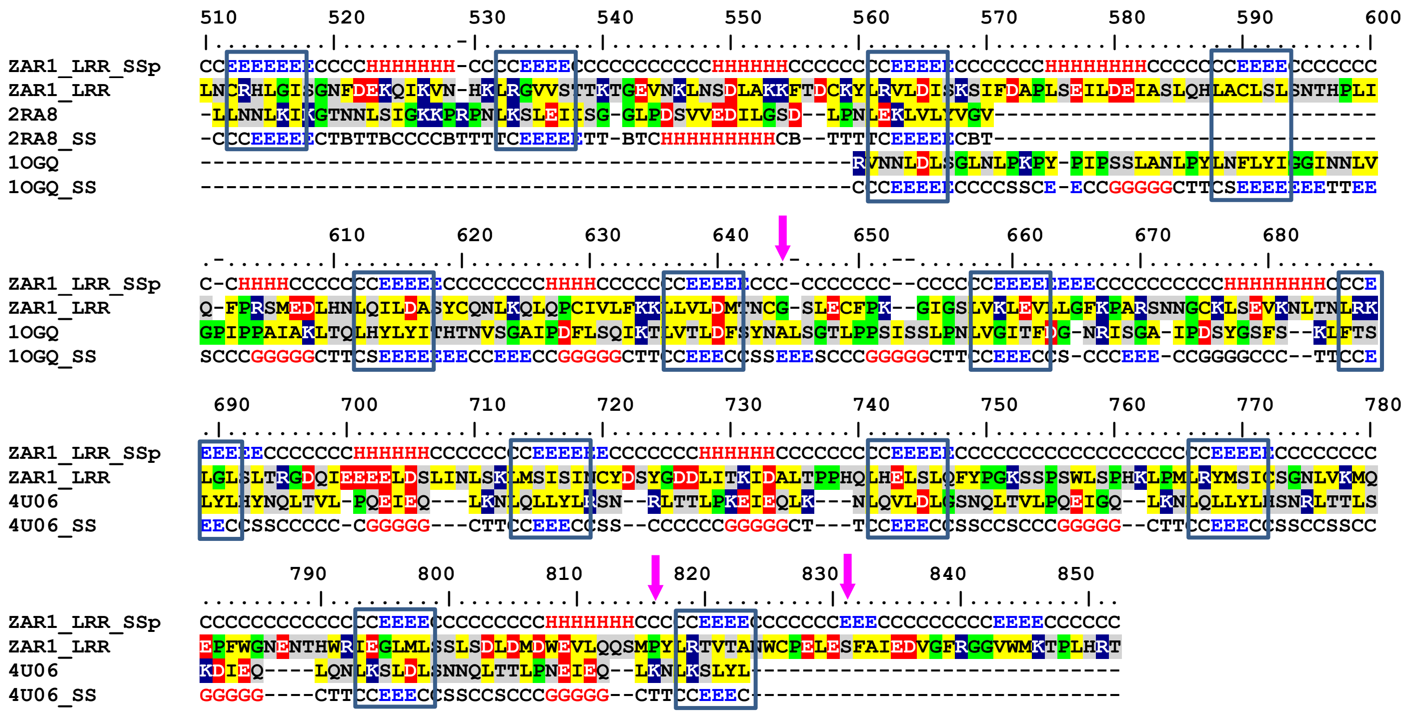

**Figure S8: ZAR1$^{LRR}$ alignments and predicted/crystal secondary structure used in modeling the LRR domain**. LRR motifs are shown in blue boxes. The location of G645E, P816Q and S831F are shown with magenta arrows. Amino acids are colored based on their properties as follows: hydrophobic (yellow), positively charged (blue), negatively charged (red), proline and glycine (green), and polar (grey).
